# CATSPERε extracellular domains are essential for sperm calcium channel assembly and activity modulation

**DOI:** 10.1101/2024.11.18.624146

**Authors:** Jae Yeon Hwang, Huafeng Wang, Jong-Nam Oh, Sarah F. Finnegan, Niels E. Skakkebaek, Jean-Ju Chung

## Abstract

The flagellar-specific Ca^2+^ channel CatSper is a multiprotein complex that is critical for successful fertilization by controlling the sperm Ca^2+^ signaling in space and time. Large extracellular domains (ECDs) of four single-pass transmembrane subunits, CATSPERβ, γ, δ, and ε, form a unique canopy structure over the pore-forming channel. However, the molecular mechanisms of canopy assembly during development and its physiological function in mature sperm remain unknown. Here, using two genetic mouse models and the biochemical isolation of a bioactive CATSPERε fragment, we report that CATSPERε ECDs are essential for assembling the CatSper canopy, and thus the entire channel complex, and for modulating CatSper function for sperm hyperactivation and fertilization. CATSPERε-deficient males are sterile because their sperm fail to develop hyperactivated motility due to the absence of the entire channel. In transgenic mice overexpressing CATSPERε with truncated ECDs in testicular germ cells, truncated CATSPERε is unable to interact with native CatSper subunits and incorporate into the complex, thus failing to rescue the defective sperm hyperactivation and infertility of *Catspere*-null males. These findings provide insight into the underlying molecular and developmental mechanisms of CatSper complex assembly and how CatSper channels can be modulated in physiological settings and by therapeutic intervention.

## Introduction

All living organisms enclosed by the plasma membrane communicate with the extracellular environment through membrane-associated proteins on their surface, such as membrane receptors and ion channels. The extracellular domains (ECDs) of these proteins are critical for sensing and transducing external signals into the cells. Typical examples are AMPA and NMDA receptors, where ligand binding to the ECDs triggers intracellular signaling (**Chen et al., 2017; Chou et al., 2024**). ECDs of various ion channels including PIEZO1 and Ca_v_1.2, often modulate ion conductance through the pore by undergoing conformational changes, thus participating in coordinating the opening or closing of the pore and influencing its activated or inactivated state (**Gao et al., 2023; Xiao, 2024**). Based on the structural information and functional importance, recent studies have reported various drugs targeting these ECDs to modulate the activity of ion channels and receptors and the resulting downstream events (**Huang et al., 2024**). Notably, diazepam, which is used to treat anxiety and seizures, targets ECDs of GABA_A_ receptors for its potentiation of GABA response to suppress neuronal activity (**Zhu et al., 2018**).

In mammalian spermatozoa, various membrane proteins are housed in the flagellar membrane for their navigation to successful fertilization in the female reproductive tract (**Wachten et al., 2017; Wang et al., 2021**). Among them, the multiprotein CatSper Ca^2+^ channel complex presents the largest ECDs. Each of the four single transmembrane accessory subunits (CATSPERβ, γ, δ, and ε) interacts with one specific pore-forming subunit, forming a huge canopy structure over the heterotetrameric channel and linking the individual holo-complexes extracellularly in zigzag rows (**Lin et al., 2021; Zhao et al., 2022**). CatSper-mediated Ca^2+^ signaling is critical for nearly all events in sperm capacitation (*i.e*., the collective process of acquiring fertility), including the development of hyperactivated motility and the acrosome reaction, and thus male fertility (**Hwang and Chung, 2023**). However, our understanding of the physiological function of the CatSper canopy and the outcome of its intervention as a drug target remains largely unexplored.

Our previous study showed that CATSPERδ is required to maintain the stability of a pore-forming CATSPER1 subunit in developing male germ cells, suggesting its role in channel assembly (**Chung et al., 2011**). Indeed, recent advances in structural elucidation of the CatSper holo-complex and *in situ* higher order arrangement further support that the canopy-forming subunits contribute not only to channel complex assembly but also potentially to activity regulation (**Lin et al., 2021; Zhao et al., 2022**). Because the entire CatSper channel was absent in mature spermatozoa from *Catsperd*-null males (**Chung et al., 2011**), the detailed molecular mechanisms of assembly and the function of canopy ECDs in mature sperm remain unknown. This critical hurdle highlights the need for a different approach to further elucidate the physiological function of the canopy and individual TM subunits in mature spermatozoa. In the CatSper holo-complex, CATSPERε, paired with the CATSPER2 subunit (**Lin et al., 2021**), is located on the wing side of the CatSper zigzag rows (**Zhao et al., 2022**) in the quadrilinear Ca^2+^ signaling nanodomains (**Chung et al., 2017; Hwang et al., 2019**). Interestingly, among the CatSper genes, human mutations in CATSPERE and its pairing partner CATSPER2 have been reported more frequently (**Brown et al., 2018; Young et al., 2024**), suggesting that their structural location is potentially vulnerable to cause functional defects in the CatSper channel and thus male infertility.

Here, using genetic and molecular dissection of CATSPERε, we show the molecular mechanisms of CATSPERε ECDs in channel complex assembly during germ cell development and their physiological function in mature spermatozoa. Genetic ablation of *Catspere* impairs sperm to develop hyperactivated motility as it leads to the absence of the entire CatSper complex, resulting in male infertility. Overexpression of the canopy roof-truncated CATSPERε prevents its interaction with other CatSper canopy-forming subunits in testicular germ cells. Finally, we demonstrate that treatment of spermatozoa with purified Ig-like domain of CATSPERε ECDs during capacitation attenuates CatSper activity, impairing sperm hyperactivation and fertilization *in vitro*. These findings provide direct insights into the canopy function in CatSper complex assembly and activity regulation during sperm capacitation, and suggest a strategy to target CatSper activity to control sperm fertility.

## Results

### CATSPERε is indispensable for sperm hyperactivation and male fertility

Four single-pass transmembrane (TM) subunits (CATSPERβ, γ, δ, and ε) with their large extracellular domains (ECDs) form the canopy structure of the CatSper channel complex (**Lin et al., 2021**). The channel complexes are arranged in zigzag nanodomains along the sperm tail with CATSPERε uniquely positioned on the wing side (**Zhao et al., 2022**; **Figure 1A** and **1B**). However, the detailed roles of CATSPERε and its physiological significance are currently unknown. Like other CATSPER subunits, *Catspere* is expressed exclusively in the testis (**Figure 1C; Figure S1A**), and CATSPERε is specifically localized to the flagellar membrane in the principal piece of human and mouse spermatozoa (**Chung et al., 2017**; **Figure 1D** and **1E**). To understand the physiological significance of CATSPERε, we generated *Catspere*-null (*Catspere*^-/-^) mice using CRISPR/Cas9 genome editing (**Figure S2A and S2B**). By introducing two guide RNAs targeting the 1^st^ and 2^nd^ exons, respectively, we obtained a null allele predicted to produce a frame-shifted protein after the 3^rd^ amino acid. CATSPERε is absent in the testis and epididymal spermatozoa of *Catspere^-/-^* males (**Figure 1E** and **F; Figure S1B; Figure S2C**). *Catspere^-/-^* mice show no gross abnormalities in survival, appearance or behavior, and null females are fertile. However, despite normal sperm production (**Figure S2D**), *Catspere*^-/-^ null males are completely infertile (**Figure 1G and S2E**), as are genetic disruptions of genes encoding other TM CatSper subunits (**Ren et al., 2001; Quill et al., 2003; Qi et al., 2007; Chung et al., 2011; Huang et al., 2023; Hwang and Chung, 2023**). To understand how CATSPERε deficiency leads to male infertility, we examined sperm motility parameters using a Computer-Assisted Semen Analyzer (CASA). Curvilinear velocity (VCL) and lateral head displacement (ALH) are significantly lower in sperm lacking CATSPERε after induction of capacitation (**Figure S2F**). Flagellar waveform analysis also revealed that *Catspere*^-/-^ sperm fail to increase the maximum angle of the primary curvature in the midpiece (α-angle) and beat in an overall smaller envelope (**Figure 1H** and **1I**), all indicative of defective sperm hyperactivation. These results demonstrate that CATSPERε deficiency impairs hyperactivated motility, resulting in male infertility.

**Figure 1.**
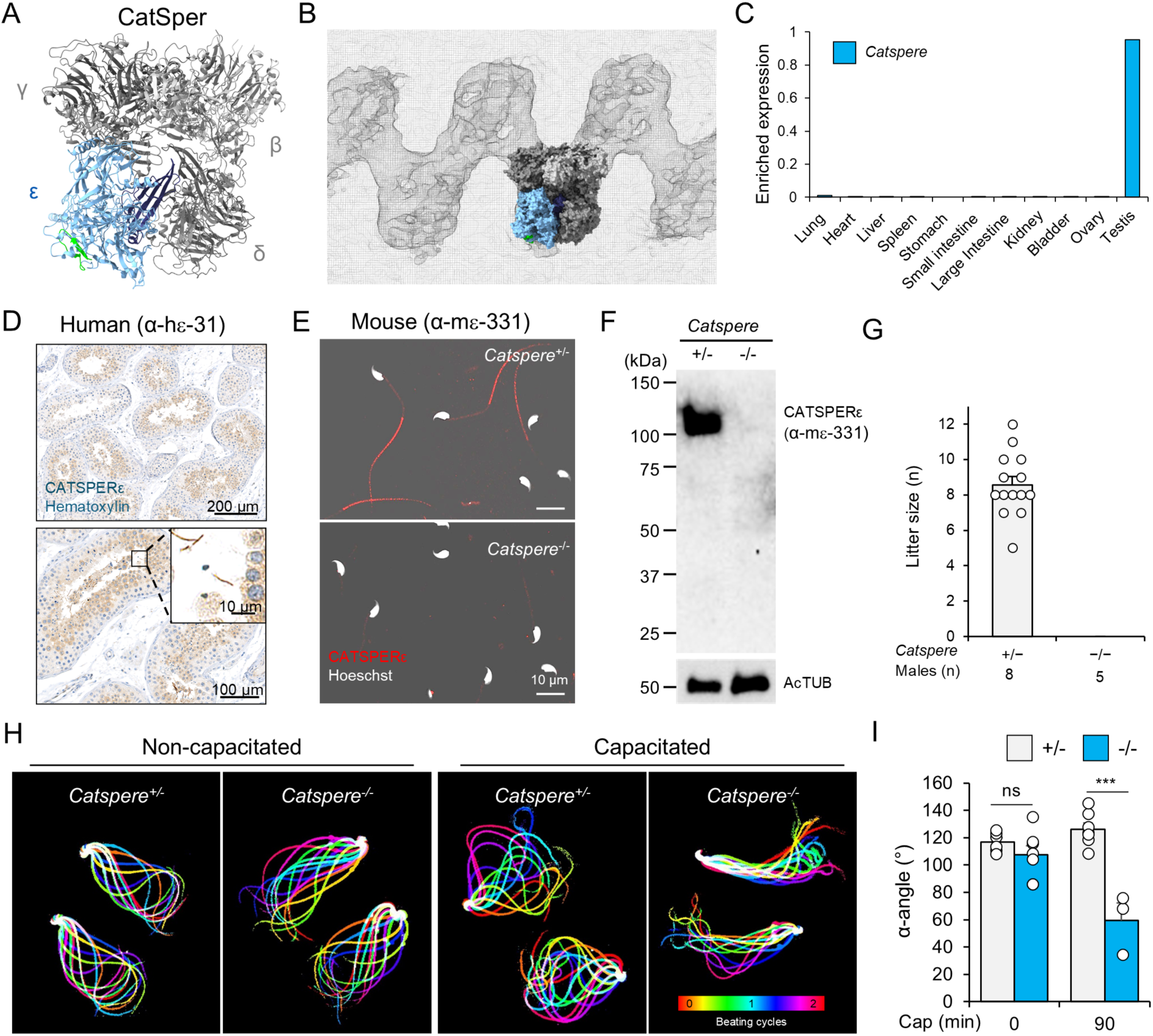
The transmembrane CATSPERε subunit is indispensable for sperm hyperactivation and male fertility. (A-B) Molecular position of CATSPERε within the atomic model and cryo-ET map of the mouse CatSper channel complex. Shown are top-down views of CATSPERε (blue) in the single holo CatSper complex (A, PDB: 7EEB), which was fitted into the cryo-ET map of CatSper (B, EMDB: 24210). The epitope used to raise α-CATSPERε (α-mε-331) that recognize an ECD region (**Nand et al., 2023**; related manuscript by **Clouser et al., 2024**) is colored in green and the Ig-like domain is colored in navy. (C) *Catspere* mRNA expression in mouse tissues. *Catspere* (blue bars) is specifically expressed in adult testis. (D) Immunostained CATSPERε in human testis. Human CATSPERε antibody (α-hε-31 (**Chung et al., 2017**) was used. Shown in magnified inset (*bottom*) are testicular sperm cells. (E) Detection of CATSPERε in live epididymal sperm from heterozygous (+/-; *top*) and knockout (-/-; *bottom*) mice. Shown are merged images of fluorescence and corresponding DIC field images. DNA is stained with Hoechst. (F) Immunoblotting of CATSPERε in epididymal sperm from *Catspere*^+/-^ and *Catspere*^-/-^ (-/-) mice. Acetylated tubulin (AcTUB) is a loading control. (G) Litter sizes from fertile females mated with *Catspere*^+/-^ or *Catspere*^-/-^ males. Individual litter sizes are marked. (H) Flagellar waveform of the epididymal sperm from *Catspere*^+/-^ and *Catspere*^-/-^ mice before (*left*) and after (*right*) inducing capacitation *in vitro*. Flagellar movement of the head-tethered sperm were recorded at 200 fps speed. The tail movements for two beating cycles are overlaid to show the waveform color-coded in time. (I) Maximum angles of the primary flagellar curvature (α-angle) of the epididymal sperm. α-angles of *Catspere*^+/-^ (+/-, gray bars) and *Catspere*^-/-^ (-/-, blue bars) sperm were measured before (0 min; +/-, 117.0° ± 2.9°; -/-, 107.6° ± 6.7°) and after (90 min; +/-, 126.2° ± 4.6°; -/-, 59.6° ± 12.8°) inducing capacitation (cap). ns, non-significant; ***p<0.001. Data are represented as mean ± SEM (G and I). *See also* Figures S1 and S2.

### *Catspere*-null sperm lack the entire CatSper channel, resulting in impaired Ca^2+^ signaling

To understand how genetic ablation of *Catspere* impairs sperm to develop hyperactivated motility, we examined whether the CatSper channel is present and functional in *Catspere*^-/-^ sperm (**Figure 2**). All CatSper subunits examined are absent in *Catspere*^-/-^ sperm (**Figure 2A** and **B**), suggesting the lack of the entire CatSper channel, consistent with the observation in *Catsperd*^-/-^ sperm (**Chung et al., 2011**). *Catspere*^-/-^ sperm prematurely enhance capacitation-mediated tyrosine phosphorylation compared with wild-type (WT) sperm (**Figure 2C and Figure S2G**), indicating impaired Ca^2+^ influx during capacitation (**Chung et al., 2014**). To directly test for defective Ca^2+^ influx through the CatSper channel, we performed electrophysiological recordings to measure the CatSper current (*I_Catser_*) in *Catspere*^-/-^ spermatozoa (**Figure 2D-F**). When elicited under voltage ramp or step protocols, *I_Catsper_* is undetectable in *Catspere*^-/-^ spermatozoa, indicating the absence of functional CatSper channels in spermatozoa. These results demonstrate that *Catspere*^-/-^ spermatozoa lack the CatSper channel and therefore fail to transduce Ca^2+^ signals to develop hyperactivated motility during capacitation.

**Figure 2.**
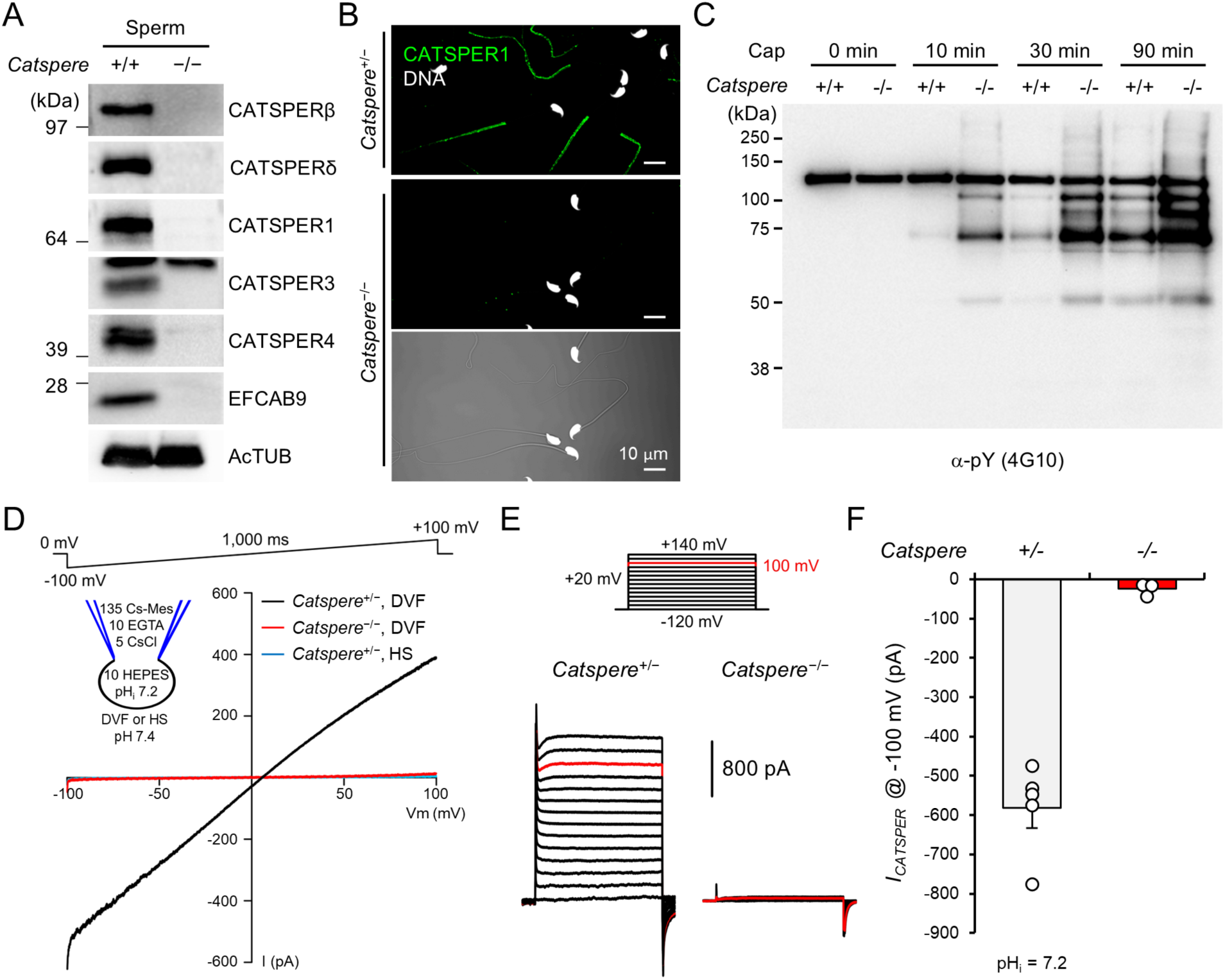
*Catspere*-null spermatozoa lack functional CatSper channel. (A) Immunoblotting of CatSper subunits in WT (+/+) and *Catspere*-null (-/-) epididymal sperm. Acetylated tubulin (AcTUB) is a loading control. (B) Confocal images of immunostained CATSPER1 in *Catspere*^+/-^ (*top*) and *Catspere*^-/-^ (*middle* and *bottom*) epididymal sperm. Fluorescence and corresponding DIC images are shown for *CatSpere*^-/-^ sperm (*bottom*). Hoechst is used for counter staining DNA. (C) Immunoblotting of protein tyrosine phosphorylation (pY) in WT (+/+) and *Catspere*-null (-/-) sperm. Time-course development of pY was examined for 90 min incubation of cauda epididymal sperm under capacitation (Cap) conditions. (D and E) Representative traces of CATSPER current (*I_CatSper_*) of corpus epididymal sperm from *Catspere*^+/-^ and *Catspere*^-/-^ males. *I_CatSper_* traces were elicited by voltage-ramp protocol from −100 mV to +100 mV range with 0 mV holding potential (D) or by step protocol from −120 mV to +140 mV ranges with 20 mV increments (E). A cartoon in panel D represents pH and ion composition (mM) in pipette solution and used bath solution for *I_CatSper_* measurement by voltage-ramp and step protocols. Divalent free (DVF) solution or HEPES-buffered saline (HS) were used for bath solutions. (F) Inward *I_CATSPER_* measured from *Catspere*^+/-^ (gray, n=5) and *Catspere*^-/-^ (blue, n=3) corpus sperm at −100 mV. Circles indicate the current from individual spermatozoa. Data are shown to mean ± SEM. *See also* Figure S2.

### Truncated CATSPERε lacking the extracellular canopy roof does not interfere with the assembly of native full-length CATSPERε into the CatSper complex

The function of the unique large extracellular canopy above the pore-forming channel remains unknown. It could serve as a binding site for natural ligands, as a linker to multimerize the channels into a linear cooperative working unit, and/or as a chaperone to assemble the pore-forming tetramer in the specific counterclockwise order (**Lin et al., 2021; Zhao et al., 2022**). Notably, sequence comparison analysis revealed that the C-terminal Ig-like domain and the stem region of CATSPERε are highly homologous to those of CATSPERδ (**Chung et al., 2017; Figure S3A**). In contrast to CATSPERβ and γ, which mediate intra- and inter-dimer interactions, respectively, CATSPERε and δ do not participate in the supramolecular interactions in the higher-order arrangement of the CatSper complexes (**Zhao et al., 2022**). Thus, we hypothesized that this homology at the level of the canopy pole (*i.e.*, the stem and TM region only without the canopy roof) is more indicative of its function in channel assembly. To test this idea at the canopy pole and to further delineate the specific role of the canopy roof, we generated a transgenic mouse line overexpressing the roof-truncated, pole-retaining CATSPERε (*i.e*., truncated CATSPERε) specifically in the testis using the calmegin promoter (**Watanabe et al., 1995; Ikawa et al., 2001**; **Figure 3A and Figure S3A**).

**Figure 3.**
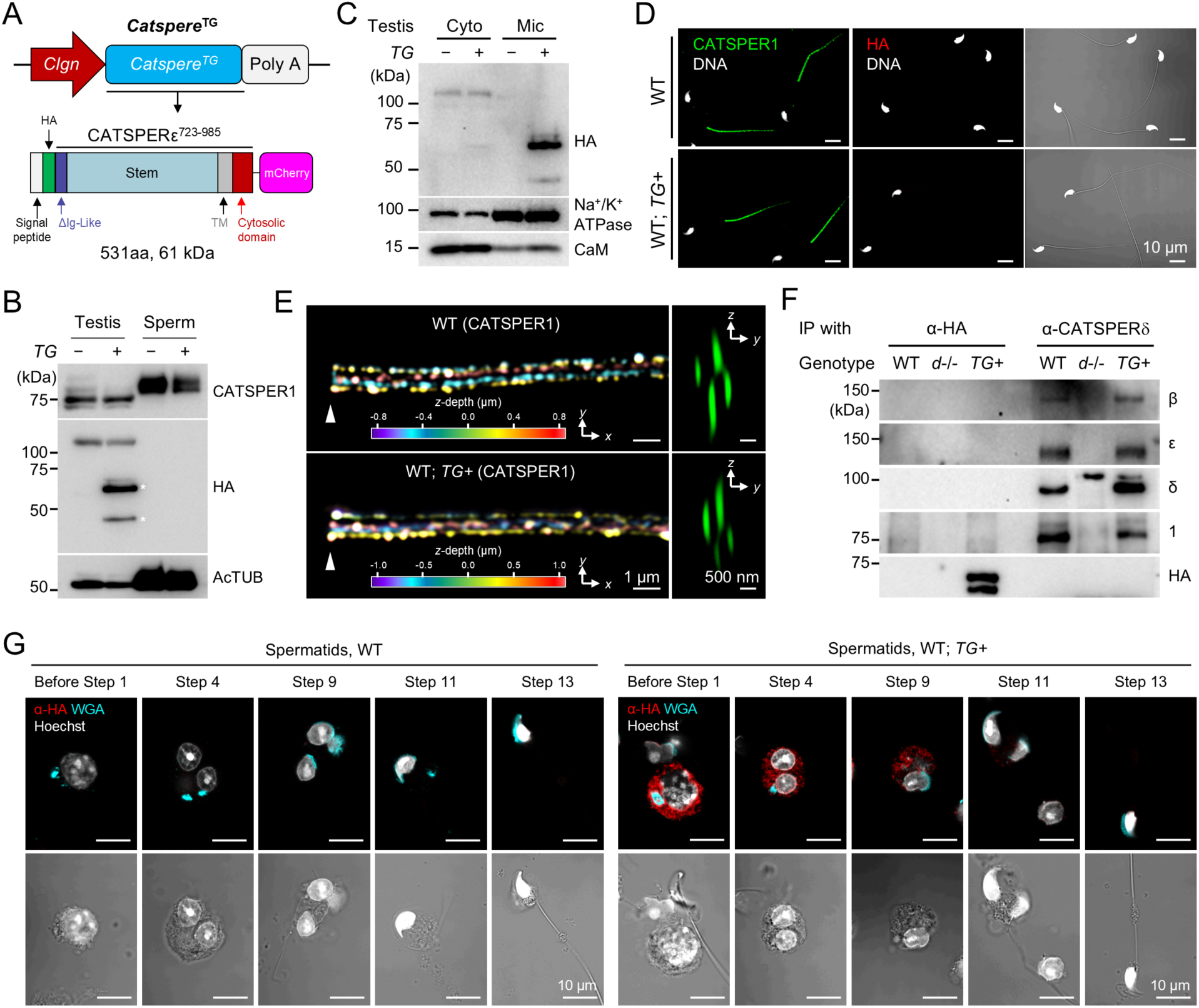
Truncation of the extracellular domain fails CATSPERε to localize at the sperm flagella. (A) A schematic diagram for *Catspere* transgene (*Catspere*^TG^) encoding extracellular domain (ECD) truncated CATSPERε in mice. The transgene encodes the CATSPERδ-homologous region of CATSPERε fused to its native signal peptide and HA at N-terminus and mCherry at C-terminus (∼61 kDa, *bottom*), respectively. *Clgn* promoter is for transgene expression specific to testicular germ cells (*top*). (B) Immunoblotting of the truncated CATSPERε in testis and epididymal sperm from *Catspere*^TG+^ (*TG+*) mice. Asterisks (*) indicate truncated CATSPERε proteins. Acetylated tubulin (AcTUB) is used as a loading control. (C) Immunoblotting of the truncated CATSPERε in cytosol and microsome fraction of testis from WT; *TG+* males. Na^+^/K^+^ ATPase and Calmodulin (CaM) serve as controls for microsome and cytosol fraction, respectively, and for loading. (D) Confocal images of immunostained CATSPER1 and HA in epididymal sperm from WT (*top*) and WT with the transgene (WT; *TG+*, *bottom*) male mice. Merged are fluorescence images of sperm immunostained with α-HA and corresponding DIC images (*right*). (E) 3D structural illumination microscopy images of immunostained CATSPER1 in sperm from WT (*top*) and WT; *TG+* (*bottom*) mice. Colors in *x-y* projection indicate *z*-depth information (*left*). *y-z* projection represents cross-section images (*right*). DNA was counter-stained with Hoechst. (F) Co-immunoprecipitation of native CATSPER subunits and truncated CATSPERε in WT, *Catsperd*-null (*d-/-*), and WT; *Catspere^TG^* (*TG+*) testis. Native CATSPER subunits – β, δ, ε, and 1 – and truncated CATSPERε are examined in HA and CATSPERδ immunocomplexes reciprocally. (G) Confocal images of truncated CATSPERε immunostained with α-HA in developing spermatids from WT (*left*) and WT; *TG+* (*right*) males. Shown are fluorescence (*top*) and corresponding DIC images merged with fluorescence for DNA (*bottom*). Wheat germ agglutinin (WGA) and Hoechst were used to stain sugar residues and DNA, respectively. The truncated CATSPERε is probed by α-HA (B, C, D, F, and G). *See also* Figure S3.

The truncated CATSPERε is expressed in testis and is enriched in the microsomal fraction, indicating its membrane association (**Figure 3B** and **3C**). However, the truncated protein is not detected in epididymal spermatozoa (**Figure 3B** and **3D**). At the same time, the presence of the truncated CATSPERε in testis did not alter the typical quadrilinear localization of the CatSper channel in the principal piece of mature sperm tail (**Figure 3D** and **3E**). These results demonstrate that only the CatSper channel complexes assembled without the truncated CATSPERε are properly trafficked to the flagellar membrane. To test this idea, we first determined whether the truncated CATSPERε is complexed with other CatSper subunits. We performed co-immunoprecipitation using testis lysate (**Figure 3F**). Native full-length CATSPERε and other examined CatSper subunits are detected in the CATSPERδ immunocomplex in the testis, independent of the transgene expression encoding the truncated CATSPERε. However, the immunocomplex of the truncated CATSPERε protein with anti-HA antibody does not contain any CatSper subunits in the transgenic animal. This result demonstrates that, contrary to our hypothesis, the canopy pole is not sufficient to assemble the canopy structure but the roof-forming ECDs of CATSPERε are also necessary for the assembly of the CatSper channel holo-complex. Without this region, the truncated CATSPERε is retained in the cell body and progressively depleted in developing spermatids (**Figure 3G**), supporting its absence in mature spermatozoa (**Figure 3B** and **3D**) and minimal effect on the assembly of native CATSPERε into the CatSper complex (**Figure 3F**). Accordingly, spermatozoa from the animal carrying both WT and transgenic alleles (WT; *TG+*) develop normal CatSper-mediated hyperactivated motility with normal tyrosine phosphorylation after inducing capacitation (**Figure S3B and S3C**). Also, protein levels of the native CatSper subunits are not altered in WT; *TG+* spermatozoa (**Figure S3D and S3E**).

### The canopy roof-truncated CATSPERε fails to form heterotetrameric auxiliary complex in *Catspere*-knockout mice

In the absence of full-length CATSPERε, the truncated CATSPERε could replace the assembly function of native CATSPERε by incorporation into the CatSper complex, while not being dominantly or equally in the presence of full-length CATSPERε (**Figure 3**). To test this idea, we crossed the transgenic mice in the background of *Catspere^-/-^* mice to express only the transgene (*Catspere*^-/-^; *TG+*; **Figure 4A**). The truncated CATSPERε proteins are well expressed in the testis but not detected in the epididymal spermatozoa as observed in the spermatozoa of WT; *TG+* males. CATSPERδ co-immunoprecipitation with testis lysate shows that truncated CATSPERε is not in complex with any of the native CatSper subunits examined, not even with the neighboring CATSPERδ or γ (**Figure 4B**). This result demonstrates that the ECDs corresponding to the canopy roof of CATSPERε are indispensable for the assembly of all four auxiliary subunits into a fourfold complex via its interaction with CATSPERδ and γ.

**Figure 4.**
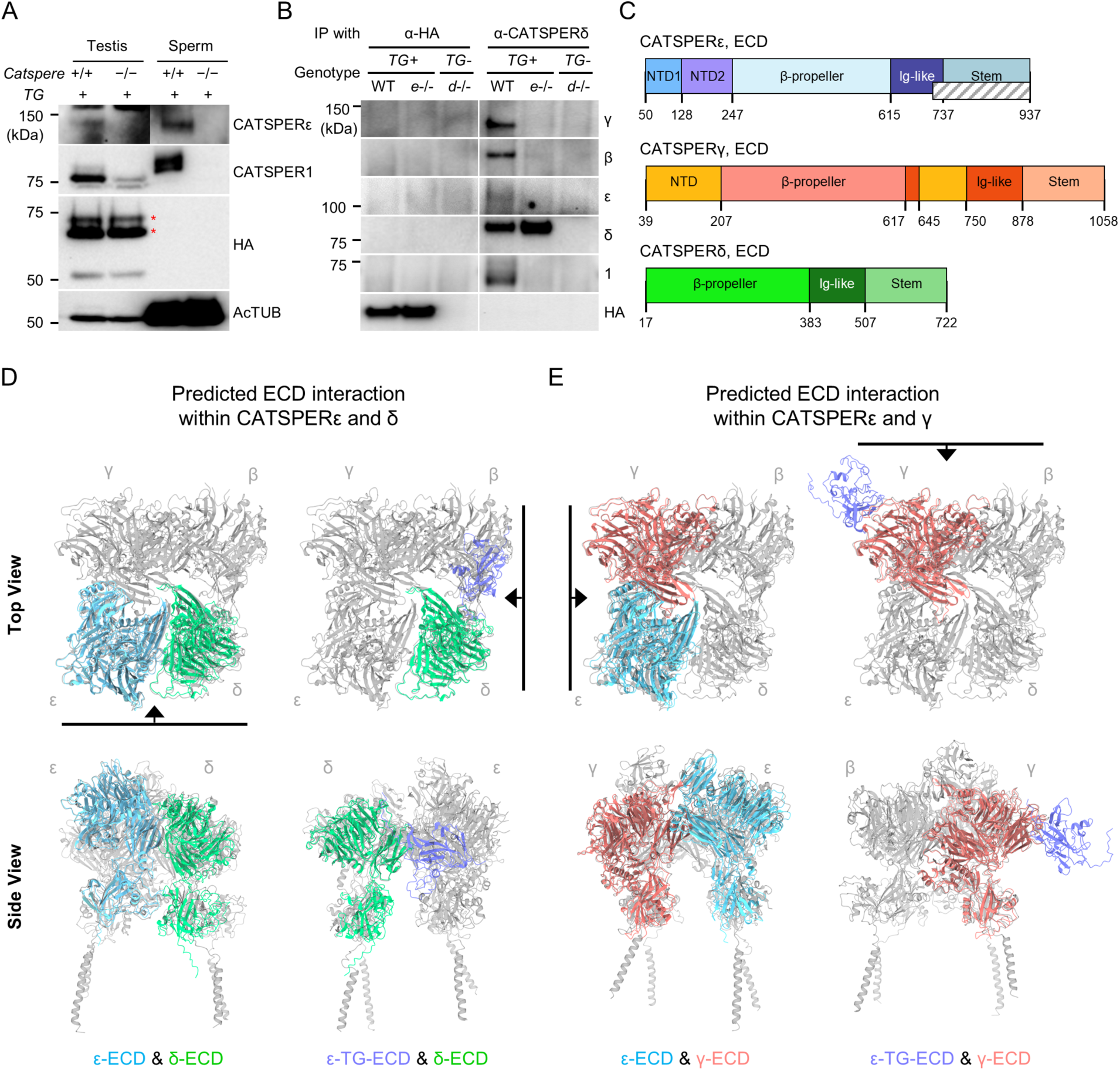
Protein-protein interaction of CATSPERε canopy with neighboring ancillary subunits is essential to assemble the entire CatSper channel complex. (A) Immunoblotting of truncated CATSPERε in testis and sperm from WT (+/+) and *Catspere*-null (-/-) males with the transgene (*TG*). Acetylated tubulin (AcTUB) is a loading control. Asterisks (*) indicate the truncated CATSPERε expressed from the transgene. (B) Co-immunoprecipitation of the truncated CATSPERε and CATSPER subunits in testis from WT and *Catspere*-null (*e*-/-) mice carrying the transgene (*TG*). α-HA was used to detect the truncated CATSPERε (A and B). *Catsperd*-null testis (*d*-/-) is used as negative control. (C) Organization of CATSPERε, γ, and δ extracellular domains (ECDs). Domain boundaries of each subunit are labeled. A gray sash in CATSPERε indicates ECD region remaining in the truncated CATSPERε. (D and E) Predicted interaction of the truncated CATSPERε and CATSPERδ (D) but not with CATSPERγ (E). Shown are AlphaFold Multimer-predicted interaction of the whole (*left*) or truncated (*right*) CATSPERε ECD with whole ECDs of CATSPERδ (D) or CATSPERγ (E). Modeled structures for interacting two ECDs are fitted to the reference canopy structure of the CATSPER channel colored in gray (PDB: 7EEB). Black lines marked in top view structures (*top*) indicate the orientation for the corresponding side views (*bottom*).

Next, to understand how the CATSPERε truncation affects the assembly of the CatSper canopy and the entire CatSper complex, we analyzed the AlphaFold predicted interactions of truncated CATSPERε with δ or γ (**Figure 4C-E**). In the predicted models, the intact CATSPERε ECDs interact normally with both δ and γ, and the predicted ε-δ and ε-γ dimers fit well into the atomic model of the CatSper canopy (CATSPERγ-β-δ-ε from PDB:7EEB) (**Figure 4D** and **E**). However, the truncated CATSPERε ECDs (*i.e.*, C-terminal Ig-like domain and stem region without TM domain) are predicted to interact abnormally with these neighboring subunits, and the aberrant dimers do not fit the reported structure. Thus, the truncated roofless partial ECDs are likely to have altered their interaction with other canopy-forming auxiliary subunits, resulting in unsuccessful canopy assembly and canopy interactions with the pore-forming channel (CATSPER1-4-3-2).

### The truncated CATSPERε fails to rescue impaired sperm hyperactivation by CATSPERε deficiency

Truncation of the canopy roof alters the interaction of CATSPERε with δ and γ, impairing ε incorporation into the canopy and thus into the entire CatSper channel complex (**Figure 3** and **4**). In addition, the truncated CATSPERε appears to have little effect on the assembly and trafficking of the CatSper channel complex in the presence of native full-length CATSPERε in sperm from both WT and *Catspere*^+/-^ males (**Figure S3, S4A-S4E**). Thus, we predict that *Catspere*^-/-;^ *TG*+ spermatozoa will lack a functional CatSper channel and will not rescue the physiological defects of *Catspere*^-/-^ animals.

Indeed, all CatSper subunits examined are absent in *Catspere*^-/-^; *TG+* spermatozoa (**Figure 5A** and **5B, Figure S4F**). Capacitated *Catspere*^-/-^; *TG+* spermatozoa aberrantly potentiate tyrosine phosphorylation (**Figure S4G**), indicating impaired CatSper-mediated Ca^2+^ signaling. Electrophysiological recording shows that *I_Catsper_* is not detectable in *Catspere*^-/-^; *TG+* spermatozoa (**Figure 5C-E**), which is not significantly different from that recorded in *Catspere*^-/-^ spermatozoa (**Figure 2D-F)**. Although *Catspere*^-/-^; *TG+* males produce a comparable number of spermatozoa as heterozygous males (**Figure S4H**), they are infertile (**Figure 5F and S4I**) with defective sperm hyperactivation (**Figure 5G and S4J**) like *Catspere^-/-^* males (**Figure 1**). All these results demonstrate that *Catspere*^-/-^; *TG+* sperm lack the CatSper channel and that truncated CATSPERε cannot rescue male infertility caused by CATSPERε deficiency.

**Figure 5.**
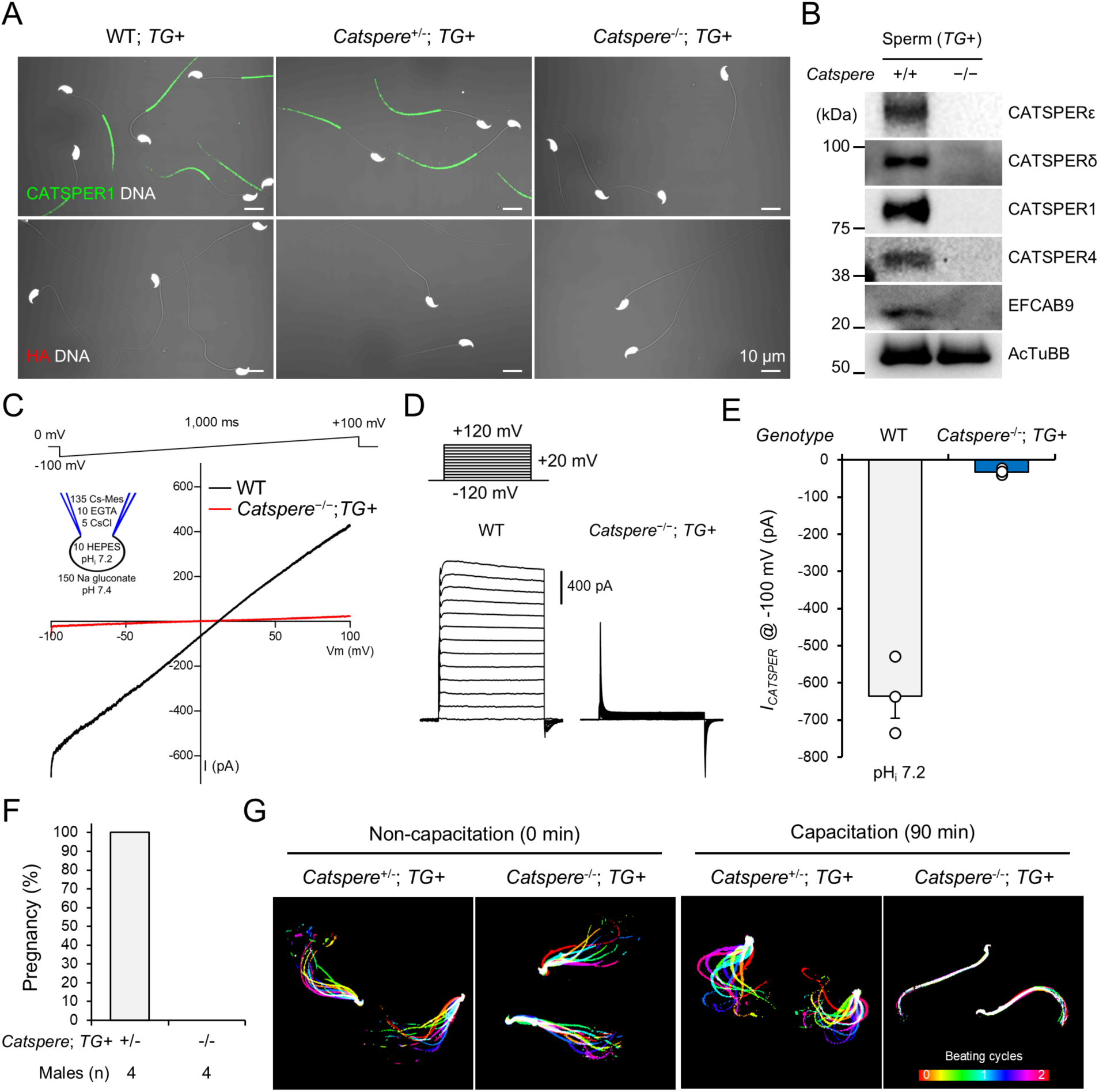
ECD-truncated CATSPERε fails to rescue impaired sperm hyperactivation and male infertility in *Catspere*-null mice. (A) Confocal images of immunostained CATSPER1 and HA in epididymal sperm from WT, *Catspere*^+/-^, and *Catspere*^-/-^ mice crossed with the transgenic (*TG*) mice. Shown are merged views of fluorescence and corresponding DIC images. Hoechst is used to counterstain DNA. (B) Immunoblotting of CATSPER subunits in epididymal sperm from WT and *Catspere*-null (*Catspere-/-*) males with the transgene (*TG*). Acetylated tubulin (AcTUB) is used as loading control. (C and D) CatSper current (*I_CATSPER_*) from sperm from *Catspere*-null mice expressing the transgene (*Catspere*^-/-^; *TG+*) and WT sperm. Shown are representative *I_CATSPER_* traces elicited by ramp (C, −100 mV to +100 mV) and step (D, −120 mV to +140 mV, 20 mM increment) protocols. Holding potential for voltage-ramp protocol and step protocol was 0 mV and −120 mV, respectively. A cartoon in panel C represents ion composition (mM) and pH in bath and pipette solution. (E) Comparison of the inward *I_CATSPER_* of corpus WT sperm (gray bar, N=3) and *Catspere*-null sperm with the transgene (*Catspere*^-/-^; *TG+*, blue bar, N=3) recorded at −100 mV. Circles indicate *I_CATSPER_* measured from individual sperm. Data are represented to mean ± SEM. (F) Pregnancy rate of WT females mated with *Catspere*^+/-^ and *Catspere*^-/-^ males carrying the transgene (*Cse*^+/-^; *TG+* and *Cse*^-/-^; *TG+*, respectively). (G) Flagellar waveform of sperm from *Catspere*^+/-^ and *Catspere*^-/-^ mice expressing the transgene (*TG+*). Tail movements of head-tethered spermatozoa were recorded at 200 fps before (non-capacitated, 0 min; *left*) and after (capacitated, 90 min; *right*) incubation under capacitating conditions. Images for two beating cycles are overlaid and color-coded in time. *See also* Figure S4.

### Ig-like domain of CATSPERε is critical in modulating CatSper activity and sperm hyperactivation

Our results using two *Catspere* genetic models provide strong evidence that the CATSPERε ECD regions that form the canopy roof are critical for their interaction with neighboring subunits to assemble the canopy and thus the CatSper channel holo-complex in developing male germ cells (**Figures 3, 4, and 5**). However, it remains to be determined whether and to what extent the canopy functions physiologically in mature spermatozoa, where the channel complexes are arranged in a higher order (**Zhao et al., 2022**).

The Ig-like domain, which is present in all four canopy-forming subunits, mediates extensive interactions between them at the center of canopy roof, whereas the stem and transmembrane domains do not (**Lin et al., 2021**). Consistently, our modeling also predicts that among the individual ECDs of CATSPERε (*i.e.,* NTD, BPD, and Ig-like), only the Ig-like domain is predicted to interact with the δ and γ ECDs at the canopy roof (**Figure S5A**), suggesting its importance in stabilizing the canopy structure. Furthermore, our preliminary solubility test of the serially deleted recombinant CATSPERε mutant proteins showed that the Ig-like domain (CATSPERε^Ig-like^; **Figure 6A**) is more soluble than other parts of CATSPERε (**Figure S5B and S5C**). Therefore, we purified recombinant mCherry-tagged CATSPERε^Ig-like^ expressed in CHO cells (**Figure 6B**) to test whether it could affect sperm physiology when administered exogenously.

**Figure 6.**
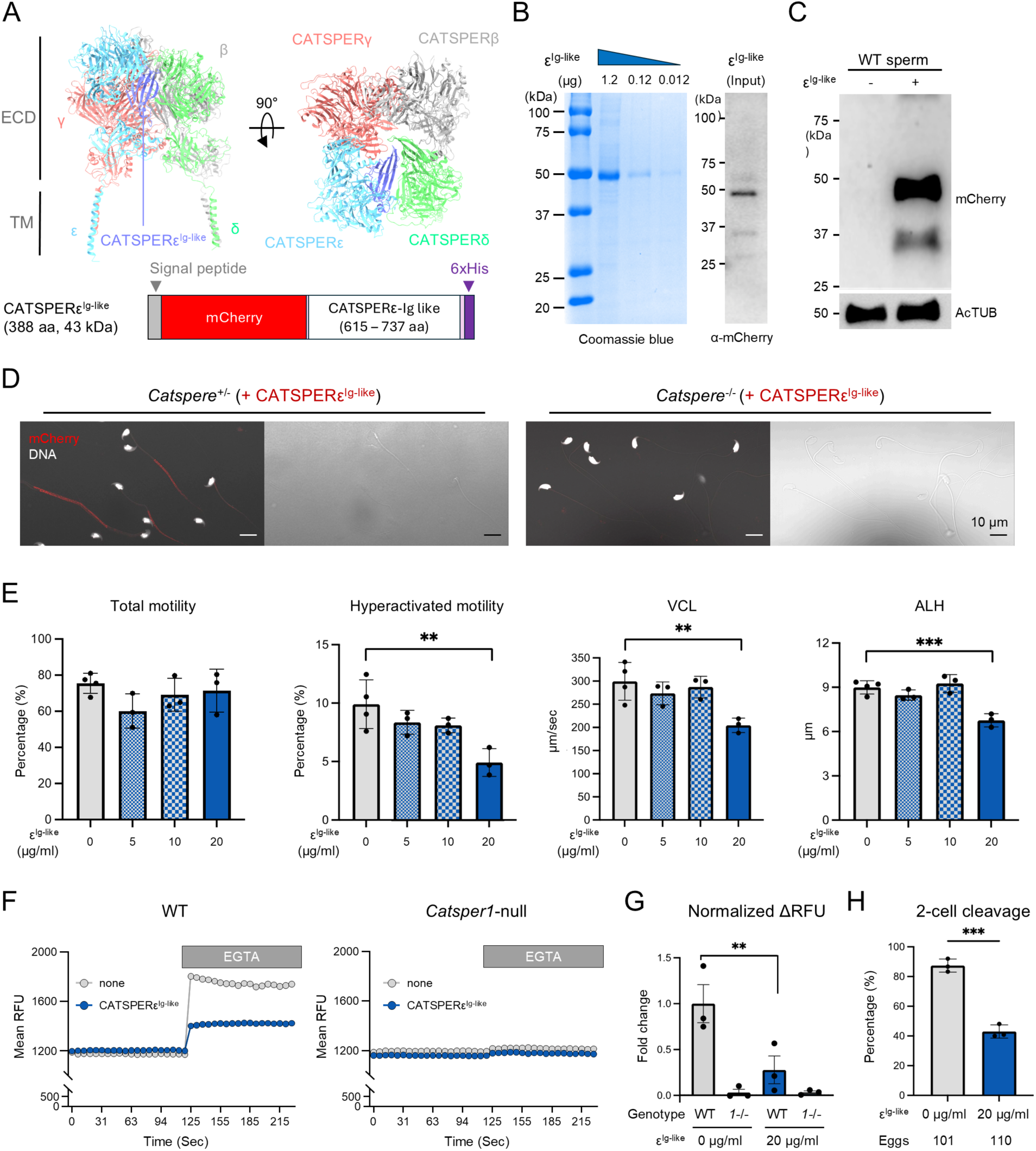
Administration of recombinant CATSPERε Ig-like domain protein attenuates CATSPER function and sperm fertility *in vitro*. (A) Location of CATSPERε Ig-like domain in the canopy structure. *Top*, shown left is the atomic model of CATSPERε, γ, δ, and β that form the canopy (from PDB: 7EEB). Shown right is top view of the canopy structure. Ig-like domain of CATSPERε (CATSPERε^Ig-like^) is highlighted in light purple. *Bottom,* a diagram of recombinant CATSPERε^Ig-like^ protein fused with mCherry and 6xHis tag. The signal peptide of mouse IgG heavy chain (gray) was added for secretion of the recombinant protein expressed in CHO cells. (B) Purified recombinant CATSPERε Ig-like domain. Shown are Coomassie blue stained (*left*) and immunoblotted (*right*) recombinant CATSPERε^Ig-like^ for purity and specificity validation. CATSPERε^Ig-like^ was probed by anti-mCherry antibody. (C and D) CATSPER-specific CATSPERε^Ig-like^ binding to epididymal sperm. Sperm cells were incubated with recombinant CATSPERε^Ig-like^ (ε^Ig-like^) proteins under capacitating conditions and the protein bound to the sperm surface are detected by immunoblotting (C) or immunostaining (D) by anti-mCherry after washing unbound proteins. Detection of recombinant CATSPERε^Ig-like^ protein added to *Catsper*^+/-^ (*left*) and *Catsper*^-/-^ (*right*) sperm. Immunofluorescence images and corresponding DIC images are shown (D). CATSPERε^Ig-like^ was detected by anti-mCherry. (E) CASA analysis of sperm motility. Total motility, hyperactivated motility, curvilinear velocity (VCL), and amplitude of lateral head (ALH) were measured from sperm after incubating under capacitating conditions with or without CATSPERε^Ig-like^. **p<0.01 and ***p<0.001. Data are represented as mean ± SD. (F and G) CATSPER activity assay by chelation-based Em changes. Live WT (*left*) and *Catsper1*-null (*right*) sperm capacitated with (gray) or without (blue) CATSPERε^Ig-like^ are loaded with ∼1 uM DiSC_3_(5) dye for Em imaging (F). Relative fluorescent unit changes (ΔRFU) after EGTA treatment are normalized and the change in WT sperm not incubated with CATSPERε^Ig-like^ was set to 1 (G). **p<0.01. N=3. Data are represented as mean ± SEM. (H) 2-cell cleavage rate of *in vitro* fertilized eggs with WT sperm capacitated with (gray) or without (blue) recombinant CATSPERε^Ig-like^. ***p<0.001. N=3. Data are represented as mean ± SD. CSε^Ig-like^, CATSPERε^Ig-like^ (E, F, G, and H).

We found that CATSPERε^Ig-like^ remained stably bound to live spermatozoa in a CatSper-dependent manner when the proteins were added to sperm incubation under capacitating conditions (**Figure 6C** and **D**). These results demonstrate that the soluble CATSPERε^Ig-like^ fragments can directly interact or intercalate with the canopy roof of the native CatSper channel complex. Recombinant CATSPERε^Ig-like^ treatment reduces motility parameters associated with sperm hyperactivation dose-dependently without affecting total motility (**Figure 6E**), suggesting reduced CatSper activity. Therefore, we performed a spectrofluometeric sperm membrane potential (Em) assay on sperm loaded with DiSC_3_(5) (**Luque et al., 2023**). CATSPERε^Ig-like^ application during sperm capacitation blocks the previously reported CatSper-specific, EGTA-induced Em depolarization by more than 70% (**Figure 6F** and **G**). Under the same conditions, CATSPERε^Ig-like^ treated spermatozoa inhibited fertilization rate of COC eggs *in vitro* by ∼ 50% (**Figure 6H**). Thus, we isolate the Ig-like domain of CATSPERε as a region important for modulating CatSper function. Our results strongly support that altering protein-protein interactions at the level of the CatSper canopy roof can prevent capacitation-induced opening of the CatSper pore-forming channels, thereby compromising sperm fertilization ability.

## Discussion

### CATSPERε ECDs are essential for assembly of the CatSper canopy and pore-forming channel

CatSper is a multiprotein channel complex composed of more than 10 TM subunits and 3 cytosolic components, arranged in higher-order zigzag rows (**Hwang and Chung, 2023**). All reported TM subunit deficiencies in mice result in the absence of a functional CatSper channel in mature spermatozoa, leading to impaired male fertility due to the defective hyperactivation (**Ren et al., 2001; Quill et al., 2003; Qi et al., 2007; Chung et al., 2011; Huang et al., 2023**) despite the presence of all other CatSper subunits in the testis except the target protein (**Chung et al., 2017; Hwang et al., 2019; Hwang et al., 2022; Huang et al., 2023**). Thus, assembly defects at various steps from the assembly of individual holo-complexes (composed of canopy and pore-forming channels) to their higher-order zigzag arrangement can be attributed to the absence of functional CatSper channels in mature sperm from these TM knockout males (**Chung et al., 2011; Hwang et al., 2022; Huang et al., 2023**). We have previously shown that CATSPERθ deficiency allows transient assembly of single holo-complexes, but results in defective interaction within CatSper dimers (**Huang et al., 2023**). The defective intra-dimer interaction prevents CatSper holo-complexes from exiting the cell body and trafficking to flagella of developing male germ cells (**Huang et al., 2023**). In contrast, the absence of either pore-forming or canopy-forming CatSper TM subunits fails to assemble transient CatSper holo-complexes in the testis (**Chung et al., 2011; Hwang et al., 2022**). For example, CATSPER1 deficiency leads to incomplete organization of the pore-forming complex, whereas canopy-forming subunits are complexed in male germ cells (**Chung et al., 2011; Hwang et al., 2022**). Deficiency of CATSPERδ or ε not only results in incomplete canopy assembly, but also reduces the protein levels of the pore-forming subunits, presumably due to their altered stability (**Chung et al., 2011; Figure 4**). Thus, a deficiency of either pore-forming or canopy-forming TM subunits likely impairs the assembly of the CatSper holo-complexes during spermatogenesis, resulting in the absence of a functional channel in epididymal spermatozoa. However, considering the earlier mRNA expression of canopy-forming subunits than pore-forming subunits during development and their different interaction patterns and protein levels in the testis (**Chung et al., 2011; Chung et al., 2017; Hwang et al., 2019**), a heterotetrameric canopy structure should form first, which presumably serves as a scaffold for 1:1 pairing with the pore-forming channels. Canopy roof-truncated CATSPERε cannot be incorporated into the CatSper complex (**Figure 4**), highlighting the importance of the protein interactions between individual canopy TM subunits for functional CatSper channel assembly. Therefore, ECD-mediated canopy assembly may be another checkpoint to ensure a complete CatSper holo-complex for flagellar trafficking and functional expression in mature sperm.

### Canopy structure is involved in modulating CatSper activity during sperm capacitation

Ion channel ECDs are important for the regulation of channel activity (**Wang et al., 2019; Nadezhdin et al., 2021**). Recent advances in cryo-EM techniques have revealed that ECDs undergo conformational changes during gating processes. For example, the ECD of GLIC, a proton-activated prokaryotic channel, is compacted and rotated counterclockwise by protonation, resulting in a widening of the pore region in the TM domains to evoke ion conductance (**Bharambe et al., 2024**). The protonated ECD of another pH-gating channel, TWIK1, closes the channel by sealing the top of the selectivity filter and suppresses K^+^ conductance (**Turney et al., 2022**). The ligand-mediated conformational changes of the ECDs and the associated gating processes have also been reported for several neurotransmitter receptors. AMPA subunits are twisted by ligand binding, resulting in conformational changes of the gating ring at the ECDs and subsequent AMPA activation (**Chen et al., 2017**). Ligand binding to other heteropentameric cys-loop family receptors, such as NMDA and glycine receptors, also induces rotation of the ECDs to propagate pore opening and channel activation (**Liu and Wang, 2023; Chou et al., 2024**). All these results show that channel activation is strongly associated with structural changes of ECDs. The current atomic structure of the CatSper holo-complex is a closed conformation resolved using the purified complex from testis and epididymis (**Lin et al., 2021**). Structural analyses revealed that the single TM domain of each canopy-forming subunit interacts with the S2 segment of the paired pore-forming subunit. In addition, their Ig-like domains interact with each other and stabilize the canopy structure at the roof just above the pore. Thus, the canopy structure connected to those of the neighboring channels might prevent the pore from opening and keep the channel in a closed conformation in non-capacitated sperm. Interestingly, exogenous addition of purified recombinant CATSPERε Ig-like domain inhibits CatSper function in sperm incubated under capacitating conditions (**Figure 6**). This result suggests that the recombinant Ig-like domain interferes with the interaction of the native CATSPERε ECD with those of the adjacent canopy-forming subunits, CATSPERγ and δ, and impairs the coordinated conformational changes of the canopy structure and/or supramolecular interaction of the zigzag arrangement during sperm capacitation. Direct structural analysis of the canopy structure in capacitated sperm will clarify this possibility and contribute to a better structural understanding of CatSper gating mechanisms during capacitation.

### CATSPERε Ig-like domain is a promising, defined drug target to intervene CatSper function

Worldwide, more than 200 million pregnancies are unintended each year (**Nickels and Yan, 2024**). Strikingly, more than 50 % of unintended pregnancies eventually end in abortion, resulting in various health and social burdens for mothers and family members (**Horvath and Schreiber, 2017; Bearak et al., 2020**). Because of the negative consequences of unintended pregnancy, there is a growing awareness and demand for improved and/or diversified contraceptive methods. Especially, men currently have very limited options such as condoms and vasectomy. To this end, the recently reported plakogloblin-SPEM1 interaction inhibitor triptonide (**Chang et al., 2021**) and soluble adenylyl cyclase inhibitor TDI-11861 (**Balbach et al., 2023**) are promising candidates for reversible male contraceptives that have been screened to target proteins specifically involved in spermiogenesis and sperm function, respectively.

CatSper is another good target validated for male fertility because of its specific expression and function in regulating sperm motility and fertility (**Mariani et al., 2023; Lee and Hwang, 2024**). However, the multi-component organization and structural complexity of CatSper has hindered the development of CatSper-targeted contraceptives. Interestingly, the large volume of ECDs and the zigzag arrangement of the CatSper channels with a 180° rotation (**Zhao et al., 2022)** suggest a uniquely high accessibility of the wing side of CATSPERε to potential biologics (*see also* the related, accompanying manuscript by **Clouser et al., 2024**). Here we show that treatment with a recombinant CATSPEREε Ig-like domain, which would disrupt the protein-protein interaction between CATSPERε and its neighbors CATSPERγ and δ within the holo-complex, inhibits CatSper and thus reduces sperm fertility (**Figure 6**). Thus, small biologics and/or small molecules that specifically bind to CATSPERε ECDs could be potential strategies to intervene in CatSper function for the development of novel non-hormonal contraceptives.

## Materials and methods

### Animals

*Catsper1 and Catsperd*-null mice (**Ren et al., 2001; Chung et al., 2011**) are maintained on a C57BL/6 background. WT C57BL/6 and CD1 mice were purchased from Charles River Laboratory. Mice were managed in accordance with the guideline approved by Yale Institute Animal Care and Use Committee (IACUC, #20079).

#### Generation of Catspere-null and Catspere-TG Mice and Genotyping

*Catspere*-null mice were generated on a C57BL/6 background using CRISPR/Cas9 system. Two guide RNAs, 5’-CGCCATGTTAGCCCGGCAGGTGG-3’ and 5’-AGAAGAACTGCAGCCTCCAGTGG-3’, in *px330* vector were injected into the fertilized eggs collected from super-ovulated C57BL/6 females after mating with males. The developing 2-Cell embryos were transplanted into pseudopregnant CD1 females and the founders’ toe were biopsied for genotyping. Truncation of the target region was examined by gDNA PCR using F2 (5’-ACTGCCCTCGTTAGCTTTTTGTCA-3’) and R1 (5’-CCTCCTTGGGCAGTTGTAGTTCA-3’) primer pair. PCR products were sanger sequenced and female founders carrying the truncated allele were backcrossed with the WT males to examine germline transmission of the allele. The *Catspere*-null mice line with 63 kb-gDNA deletion were maintained and genotyping were performed by primer pairs of F2 - R1 for the *Catspere*-null allele and F1 (5’- GCATACTAATTGCTTGGTCAAAAAC-3’) - R1 for WT allele.

Transgenic mice were generated by introducing transgene encoding the truncated CATSPERε (*Catspere-TG*) in *pClgn-mCherry* vector. Cloned *pClgn-Catspere-TG-mCherry* was linearized by restriction enzyme digestion and electroporated into the fertilized eggs collected from C57BL/6. The embryos were transplanted pseudopregnant CD1 females. Founders’ toes were clipped for gDNA extraction. Extracted gDNA were subjected for PCR with forward (5’- AAGATTTACAGGCAGTTTATTATTGAG-3’) and reverse (5’-GTCGGAGGGGAAGTTGGT-3’) primers. *Catspere-null*; *Catspere-TG* mice were generated by mating *Catspere*-TG male with WT background and *Catspere*-null females.

### Human samples (BioBank)

Human materials for the present study were collected in accordance with the principles of the Declaration of Helsinki. The regional medical research ethics committee of the Capital Region of Copenhagen approved the use of human tissues stored in the biobank at the Department of Growth and Reproduction for gene expression studies (permit number H-16019636). All patients have given informed consent to donate their residual tissue for research.

### Cell lines

HEK293T cells derived from embryonic kidney (ATCC) were cultured in DMEM (GIBCO) supplemented with 10% FBS (Thermofisher) and 1x Pen/Strep antibiotics (GIBCO) at 37 °C, 5% CO_2_ condition. The cultured cells were used to test solubility of the recombinant proteins.

### Bacterial strains

NEB 10-β bacterial strain (NEB) was used for molecular cloning.

### Mouse Sperm Preparation

Epididymal sperm from adult male mice were collected from caudal epididymis by swim-out in either M2 medium (EMD Millipore) or HEPES saline (HS medium; 135 NaCl, 5 KCl, 1 MgSO4, 2 CaCl_2_, 20 HEPES, 5 Glucose, 10 Lactic acid, and 1 Na pyruvate) adjusted to pH 7.4). Collected epididymal sperm were incubated in human tubular fluid (HTF) medium (EMD Millipore) at 2×10^6^ cells/ml concentration under the 37 °C, 5% CO_2_ condition for 90 min to induce capacitation.

### Mouse Testicular Cells Preparation

Dissociated mouse testicular cells were prepared as previously (**Hwang et al., 2021b; Hwang et al., 2022**). Briefly, seminiferous tubules in adult testes were dissociated mechanically after removing tunica albuginea. The tubules were washed with cold PBS, chopped, and minced to dissociate testicular cells. The minced tubules in PBS were filtered with 40 µm-mesh cell strainers to remove debris. Filtered testicular cells were subjected to immunostaining.

### Human Testis Section Preparation

Human testis tissues were excised from the normal, tumor-free periphery of orchiectomy specimens from four adult patients with testicular germ cell cancer. Only the specimens with tumors typical of young adults, i.e., derived from germ cell neoplasia *in situ* (GCNIS), either seminoma or nonseminoma, were included. Excised specimens were immediately fixed in GR-fixative (7.4% formaldehyde, 4% acetic acid, 2% methanol, 0.57% sodium phosphate, dibasic and 0.11% potassium phosphate, monobasic) for at least 16 hours. Samples were then dehydrated and embedded in paraffin.

### Antibodies and Reagents

Rabbit polyclonal anti-mouse CATSPER1 (**Ren et al., 2001**), 3, 4 (**Qi et al., 2007**), β (**Liu et al., 2007**), δ (**Chung et al., 2011**), EFCAB9 (**Hwang et al., 2019**), and anti-human CATSPERε (α-hε-31; **Chung et al., 2017**) were described previously.

#### Custom, in-house antibodies generated in this study

Rabbit polyclonal anti-mouse CATSPERε (α-mε-331; see also **Nand et al., 2023** and related manuscript by **Clouser et al., 2024**) was generated by immunizing rabbits with KLH-conjugated peptide (DGTVYLRTEDEFTKLDES; Open Biosystems). Rabbit polyclonal anti-mouse CATSPERγ antibody (α-mγ-957) was generated by immunizing rabbits with the KLH-conjugated peptide (RDYTEEEIFRYNSPLDTTNSLI; BioMatik). From the collected sera, the antibodies are affinity purification using AminoLink coupling resin (Pierce). A mouse monoclonal anti-CATSPER1 (clone 6D3) was established from screening hybridomas generated from mice immunized with purified N-terminus of mouse CATSPER1 (1-150 aa; GenScript) followed by Protein G affinity purification.

#### Commercially available antibodies

Mouse monoclonal antibodies against HA (clone HA-7, H3663), acetylated tubulin (clone 6-11B-1, 7451), protein tyrosine phosphorylation (clone 4G10, 05-321), and Calmodulin (05-173) were from Sigma-Aldrich. Mouse monoclonal anti-Na^+^/K^+^ ATPase was from SantaCruz (clone H-3, sc-48345). Mouse monoclonal anti-mCherry was from Novus (Novus NBP1-96752). Rabbit monoclonal anti-HA (clone C29F4, 3724) was from Cell Signaling Technology. Goat anti-mouse (115-035-003) and anti-rabbit (111-035-144) IgG conjugated with HRP were from Jackson ImmunoResearch. Veriblot for immunoblotting of the coIP was from AbCam (ab131366). Alexa 568-conjugated goat anti-mouse (A-11031) and anti-rabbit (A-11036) IgG (H+L), and Alexa 647-conjugated WGA (W32466) were from Invitrogen. Hoechst dye was from TOCRIS (5117).

All other chemicals and reagents were from Sigma Aldrich unless indicated.

### Molecular Cloning

A construct to generate *Catspere-TG* transgenic mice was cloned. Mouse *Catspere* which is sequence-homologue to the mouse *Catsperd* was subcloned into the *pClgn-mCherry* vector, gifted by Dr. Masahito Ikawa using NEBuilder® HiFi DNA Assembly Kit (NEB). Sequences encoding the native signaling peptide and HA tag are placed at the 5’ region of the *Catspere-TG* sequences. Constructs encoding fragments of the CATSPERε extracellular domains (ECDs) were subcloned into phCMV3 vector using NEBuilder® HiFi DNA Assembly Kit (NEB).

### Protein Extraction and Immunoblotting

#### Epididymal Sperm

Whole sperm proteins were extracted as described previously (Hwang et al., 2022). Collected sperm from cauda epididymis were washed with PBS followed by lysis using 2X LDS buffer for 10 minutes with vortaxing at room temperature (RT). The lysates were centrifuged at 4 °C, 18,000 x g, and supernatants were collected and denatured by incubation with 50 mM DTT at 75 °C for 10 minutes. The denatured sperm proteins were used for SDS-PAGE and immunoblotting. Primary antibodies used are: rabbit polyclonal anti-mouse CATSPERβ, CATSPERδ, CATSPER1, CATSPER3, CATSPER4, and EFCAB9 at 1 µg/ml, anti-mouse CATSPERε (1.6 µg/ml), rabbit monoclonal anti-HA (1:2,000), and mouse monoclonal anti-pTyr (1:2,000) and AcTUB (1:20,000).

#### Testis

Testis proteins were obtained as previously (Hwang et al., 2022). Testes were homogenized in 0.32M sucrose buffer using dounce homogenizer and centrifuged at 4 °C, 1,000 x g for 15 minutes to discard nucleus and debris. Collected supernatant were either lysed to 1X LDS buffer or subjected to centrifugation at 4 °C, 100,000 rpm for 1 hour to separate microsome and cytosolic fraction. Separated fractions were lysed to 2X LDS volume-equivalently. Lysed testes samples were denatured by heating at 75 °C for 2 or 10 minutes with 50 mM DTT supplement, followed by SDS-PAGE and immunoblotting. Primary antibodies for the immunoblotting were: rabbit polyclonal anti-CATSPERε (2.4 µg/ml) and CATSPER1 (1 µg/ml), rabbit monoclonal anti-HA (1:2,000), and mouse monoclonal anti-Na^+^/K^+^ ATPase (1:2,000), CaM (1:2,000), and AcTub (1:20,000). HRP-conjugated goat-anti mouse or rabbit antibodies were used for secondary antibody with 1:20,000 concentration according to the host species of the primary antibody.

#### Transfected mammalian cells

Cultured 293T cells were transfected with HA-tagged recombinant CATSPERε ECDs using linear polyethylenimine (40 kDa, Polyscience). Cells were harvested after 36 - 48 hours from transfection followed by washing with PBS two times. The transfected cells were solubilized using 1% Triton X-100 in PBS for 4 hours at 4 °C by gentle rocking and centrifuged at 4 °C, 18,000 x g for 1 hour to pellet insoluble fraction. Soluble proteins were mixed to 1x LDS buffer and insoluble pellet fractions were lysed using 2X LDS buffer volume-equivalently. Proteins in LDS sample buffer were denatured by boiling at 75 °C for 2 minutes with 50 μM dithiothreitol. Denatured samples were subjected to SDS-PAGE and immunoblotted using mouse anti-HA (1:2,000) to probe recombinant CATSPERε ECD proteins.

### Testis Co-Immunoprecipitation

Proteins in testes microsome were solubilized using PBS containing 1% Triton X-100 and cOmplete Mini, EDTA-free Protease Inhibitor Cocktail (Roche) at 4 °C for 3 hours with gentle rocking. The lysates were centrifuged at 4 °C, 18,000 x g for 1 hour and supernatants with solubilized protein were collected. Solubilized microsome proteins were incubated with SureBeads^TM^ Protein A Magenetic Beads (Bio-Rad) conjugated with rabbit polyclonal anti-CATSPERδ (0.5 µg) or mouse monoclonal anti-HA (0.5 µg) at 4 °C, overnight. The incubated magnetic beads were washed with 1% Triton X-100 in PBS and the immunocomplexes were eluted with 2X LDS buffer containing 50 mM DTT, followed by heating at 75 °C for 10 minutes. The elutes were used for SDS-PAGE and immunoblotting. Used Primary antibodies are: rabbit polyclonal anti-CATSPERγ, CATSPERδ, and CATSPER1 at 1 µg/ml, CATSPERβ (2 µg/ml), CATSPERε (2.4 µg/ml), and rabbit monoclonal anti-HA (1:1,000). Veriblot (Abcam) was used for secondary antibody with 1:200-1:500 concentration.

### Immunocytochemistry

#### Fixed cell immunocytochemistry

Collected epididymal sperm and testicular germ cells were immunostained as previously with minor modification (**Hwang et al., 2021b; Hwang et al., 2022**). Briefly, epididymal sperm and dissociated testicular cells were washed with PBS and centrifuged at 700 x g for 5 minutes or at 250 x g for 3 minutes, respectively, to attach on the glass coverslips. For testicular cells, glass coverslips were coated with poly-D-lysine before centrifugation. Attached epididymal sperm and testicular cells were fixed with 4% PFA for 10 minutes at RT and washed three times with PBS. Fixed cells were permeablized by 0.1% Triton X-100 in PBS for 10 minutes followed by blocked with 10% normal goat serum in PBS for 1 hour at RT. Blocked samples on coverslips were incubated with primary antibodies at 4 °C overnight. Primary antibodies used for the immunocytochemistry were: rabbit polyclonal anti-CATSPER1 (10 µg/ml), rabbit monoclonal anti-HA (1:200), mouse monoclonal anti-pTyr (1:100). The samples were washed with 0.1% Triton X-100 in PBS one time and PBS two times followed by incubation with goat anti-mouse or rabbit IgG conjugated with Alexa 568 (1:1,000) in blocking solution for 1 hour at RT. Testicular cells were stained with WGA conjugated with Alexa 647 (0.1 µg/ml) together with secondary antibody. Immunostained coverslips were mounted with Vectashield (Vector Laboratory) and imaged using Plan-Apochrombat 63X/1.40 oil objective lens equipped in Zeiss LSM710 Elyra P1(Invitrogen). Hoechst was used for counter staining.

#### Live cell Immunocytochemistry

Prepared sperm were incubated in M2 medium supplemented with 20 μg/mL of the anti-mouse CATSPERε at 37°C for 90 minutes. Sperm incubated with the primary antibody were washed with PBS followed by incubation with goat anti-rabbit IgG conjugated with Alexa 568 (1:1000) and Hoechst (1:500) in HS medium at 37 °C for 30 minutes. Sperm were washed with PBS and placed on the glass coverslip coated with poly D-lysine for 15 minutes followed by centrifugation at 700 x g for 5 minutes. Sperm were fixed with 4% PFA for 10 minutes at RT and washed with PBS three times. Prepared coverslips were mounted with ProLong Gold (Invitrogen) and imaged by a Plan-Apochrombat 63X/1.40 oil objective lens equipped in Zeiss LSM710 Elyra P1 (Invitrogen).

### Immunohistochemistry

Immunohistochemistry was conducted by a standard indirect peroxidase method as previously (**Nielsen et al., 2019**). Briefly, 4 µm tissue sections were subjected to heat-induced antigen retrieval using medical decloaking chamber (Biocare, Concord, CA, USA) in 0.01M citrate buffer (pH 7.4) at 110 °C for 30 min. Endogenous peroxidase was blocked with 1% H_2_O_2_ in methanol for 30 min and the sections were blocked using 0.5% skimmed milk in Tris-buffered saline (TBS) for 30 min at RT. Sections were incubated with the anti-human CATSPERε antibody (**Chung et al., 2017**) diluted 1:75 in TBS overnight at 4 °C and for 1 hour at RT in a humidified chamber. The slides incubated with the primary antibody were washed with TBS followed by incubation with anti-rabbit ImmPRESS HRP (peroxidase) antibody (Vector laboratories, CA, USA) at RT for 30 min. After incubation with secondary antibody, slides were washed with TBS and visualized using ImmPACT DAB peroxidase (HRP) substrate (Vector Laboratories). The sections were subsequently counterstained with Mayer’s hematoxylin and mounted with Aquatex® mounting medium (Merck KGaA, Germany).

### Recombinant CATSPERε^Ig-like^ proteins

Ig-like domain of the CATSPERε (CATSPERε^Ig-like^) tagged with signaling peptide sequences of mouse IgG heavy chain at N-terminus and mCherry and 6xHis at C-terminus was obtained by CHO express service (GenScript). Briefly, the recombinant protein was expressed in CHO cells and purified using HisTrap™ FF Crude (GE Health). The purified CATSPERε^Ig-like^ was subjected to SDS-PAGE followed by gel staining using Imperial^TM^ Protein Stain (Thermo Scientific^TM^) and immunoblotting with anti-mCherry to examine its purity and specificity.

### Binding test of the Recombinant CATSPERε^Ig-like^ to Epididymal Sperm

#### Immunoblotting after sperm incubation with CATSPERε^Ig-like^

Collected epididymal sperm were incubated with the purified CATSPERε^Ig-like^ and subjected to immunoblotting and live cell immunocytochemistry to test its binding to the epididymal sperm. For immunoblotting, sperm were capacitated in HTF medium supplemented with or without 20 µg/ml of the recombinant CATSPERε^Ig-like^ for 90 minutes. The capacitated sperm were washed three times with PBS to remove unbound proteins and subjected to immunoblotting as described above. Bound recombinant protein was detected by anti-mCherry (1:1,000).

#### Live cell immunocytochemistry

Collected epididymal sperm were incubated with the purified CATSPERε^Ig-like^ in capacitating conditions for 90 minutes. The sperm were washed one time with PBS followed by an hour incubation with anti-mCherry (1:1000) supplemented in mixture of 0.4% BSA-containing HEPES-buffered HTF (H-HTF; Chung et al., 2017) and immunoenhancer (Pierce) in 1:1 ratio at 37 °C. The sperm were washed two times with PBS and incubated with goat anti-mouse IgG conjugated with Alexa 568 (1:1,000) and Hoechst (1:500) in H-HTF for 30 minutes. After the incubation, sperm were washed with PBS two times and prepared on the coverslips and fixed with 4% PFA in PBS followed by washing with PBS three times for the imaging as described above.

### Structural Illumination Microscopy

Mouse epididymal sperm on coverslips were immunostained as described above and subjected to 3D structural illumination microscopy (SIM) imaging. 3D SIM imaging was performed with Zeiss LSM710 Elyra P1 using alpha Plan-APO 63X/1.40 oil objective lens (Carl Zeiss). Z-stack images was acquired with 100 nm intervals and each section was taken using images were taken using a laser at 561 nm wavelength and 5 grid rotations with a 51 nm SIM grating period. Raw images were processed and rendered using Zen 2012 SP2 software (Carl Zeiss).

### Histology Analyses

Histological analyses of the mouse tissues were performed as previously (Hwang et al., 2021b). Briefly, collected testes and epididymis were rinsed with PBS and fixed with 4% PFA in PBS at 4 °C overnight. The fixed tissues were washed with PBS and serially dehydrated by incubation in ethanol to 100%. The tissues were embedded into paraffin followed by sectioned, and the tissue sections were deparaffinized to stain with hematoxylin and eosin (H/E). The stained sections were observed using Nikon E200 microscope under 10x phase contrast objective (CFI Plan Achro 10X/0.25 pH1 WF, Nikon) and imaged by the equipped ac1300–200 μm CMOS camera (Basler AG).

### Mating Test

WT female mice were caged with *Catspere*^+/-^, *Catsper*^-/-^, *Catspere*^+/-^; *TG+*, or *Catspere*^-/-^; *TG+* males and monitored for two months to record pregnancy and litter size.

### Motility Analysis

#### Flagella Waveform Analyses

Flagellar waveform analyses were carried out as described before (**Hwang et al., 2021a; Wang et al., 2022**). Briefly, non-capacitated or capacitated epididymal sperm (2-3×10^5^ cells) were transferred to the 37°C HS and H-HTF medium, in fibronectin-coated Delta T chamber (Bioptechs). Planar flagellar movements of the head-tethered sperm were recorded for 2 seconds with 200 frame per seconds (fps) using pco.edge sCMOS camera equipped in Axio observer Z1 microscope (Carl Zeiss). Recorded images stacks were subjected to FIJI software (**Schindelin et al., 2012**) to measure beating frequency and α-angle of sperm tail (**Chung et al., 2011**), and to generate overlaid images to trace waveform of sperm flagella as previously described (**Chung et al., 2017**).

#### Computer Assisted Semen Analysis

Computer-assisted semen analysis (CASA) was carried out as previous studies (Wang et al., 2022; Hwang et al., 2022). Aliquot of non-capacitated and capacitated sperm (3.0 x 10^6^ cells/ml), which were incubated with or without recombinant CATSPERε^Ig-like^ as above, were placed in slide chamber (CellVision) and motility was examined on a 37°C stage of a Nikon E200 microscope under 10X phase contrast objective (CFI Plan Achro 10X/0.25 Ph1 BM, Nikon) equipped in Nikon E200 microscope. Images were recorded at 50 fps using CMOS video camera (Basler acA1300-200um, Basler AG, Ahrensburg, Germany) and analyzed by sperm Class Analyzer (v6.3; Microptic, Barcelona, Spain). Over 200 motile sperm were analyzed for each trial, at least 3 biological replicates were performed for each experimental group.

### In Vitro Fertilization (IVF)

4-8 weeks old female CD1 mice were injected with 10 IU of pregnant mare serum gonadotropin (PMSG, ILEX Life Science) followed by 10 IU of human chorionic gonadotropin (hCG, EMD Milipore) injection after 48 hours. 16 hours after the hCG injection, female mice were sacrificed to collect cumulus-oocyte complexes (COCs) that were placed in fresh TYH medium (**Toyoda and Chang, 1974**) covered with filtered embryo-grade mineral oil (Sigma Aldrich). Epididymal sperm were collected from WT males by swim out into TYH medium followed by capacitation in TYH medium with or without 20 μg/ml of purified CATSPERε^Ig-like^ protein for 90 minutes at 37 °C, 5% CO_2_ condition. Collected COCs were incubated for 1 hour at 37°C, 5% CO_2_ condition and supplemented with 4 x 10^5^ cells of capacitated sperm. They were incubated together for 6 hours at 37°C, 5% CO_2_ condition. After the incubation, oocytes were washed with TYH three times and transferred to fresh TYH medium. Cleaved 2-cell embryos after one day incubation were considered as fertilized eggs.

### Electrophysiology

Whole sperm patch clamping was performed as described before (**Hwang et al., 2022**). Sperm were collected from corpus epididymis and washed twice. Washed sperm were resuspended in HS medium followed by adhesion on a culture dish (Corning). Gigaohm seals were formed at the cytoplasmic droplet of motile sperm (Kirichok et al., 2006). Cells were broken through by voltage stimulation (400-600 mV, 5 ms) and suction. Whole-cell CatSper currents were recorded in the divalent-free bath solution composed of (in mM): 150 Na gluconate, 20 HEPES, and 5 Na_3_HEDTA with pH 7.4 or HS medium, pH 7.4. Intrapipette solution for recording consisted of 135 CsMeSO_3_, 5 CsCl, 10 EGTA, and 10 HEPES adjusted to pH 7.2. Data were sampled at 10 Hz and filtered at 1 kHz. The obtained current data were analyzed with Clampfit (v10.4, Axon, Gilze, Netherlands) and plotted with Grapher (v8.8, Golden Software, Inc., Gloden, Colorado).

### Chelation-Based Spectrofluorometric Sperm Em assay

Prepared WT and *Catsper1*-null sperm were capacitated in HTF with or without 20 μg/ml of CATSPERε^Ig-like^ at 2.0 x 10^6^ cells/ml concentration for 90 minutes. Incubated sperm were washed with PBS one time and resuspended into prewarmed H-HTF. 2.0 x 10^5^ cells in 100 µL of H-HTF were loaded with 10 µL of 10 µM DISC3(5) dye suspended in DMSO (American Bio, AB00435-00500). 1 x 10^5^ cells were loaded in each well of a 96-well plate and incubated for 2 minutes at 37°C. After 6 seconds of orbital shaking, fluorescence was measured for 3 minutes using Tecan Infinite 200 Pro operated by i-control software. 3.5 mM of EGTA was used to chelate free Ca^2+^ in H-HTF medium, and fluorescence levels were measured for 2 additional minutes after adding EGTA. All relative fluorescence units (RFU) were background subtracted. RFU changes (ΔRFU) were calculated by subtracting mean fluorescence intensities of EGTA-treated sperm to those of the same sperm prior to EGTA addition. Each value is taken 1 minute before and after adding EGTA respectively. ΔRFU in WT sperm capacitated without recombinant CATSPERε^Ig-like^ were set to 1 for normalization.

### Tissue and Testicular Expression analyses

Transcript data of the CatSper subunits in adult mouse and human tissues were obtained from Mouse ENCODE transcriptome data (**Yue et al., 2014**) and HPA RNA-seq normal tissues data set (**Fagerberg et al., 2014**), respectively, at NCBI (https://www.ncbi.nlm.nih.gov/gene/). Enriched tissue expression was calculated by normalizing transcript level in each tissue with sum of the transcript levels in entire tissues. Enriched tissue expression levels of the CatSper subunits are shown as heatmap using GraphPad Prism 8.0 (GraphPad Software Inc., San Diego, CA).

### Structural modeling and data analyses

A structure for mouse CatSper complex was from RCSC Protein Data Bank, PDB (https://www.rcsb.org/; 7EEB). Protein structures were rendered and visualized by iCn3D at NCBI (https://www.ncbi.nlm.nih.gov/Structure; **Wang et al., 2020**). Dimer and trimer structures for ECD domains of the CATSPER auxiliary subunits were predicted using AlphaFold-Multimer under Google Colab notebook (**Evans et al., 2021**). UCSF ChimeraX (**Meng et al., 2023**) was used to visualize atomic model of the CatSper channel (PDB: 7EEB) and to fit the predicted structures to the reference structure using matchmaker tool incorporated in ChimeraX.

### Statistical Analyses

Statistical analyses were performed by the Student’s t-test and one-way ANOVA with Dunnett’s test. Differences were considered significant at *p<0.05, **p<0.01, and ***p<0.001.

## Acknowledgement

We thank Masahito Ikawa at Osaka University for sharing *pClgn-mCherry* vector and Katerina Politi at department of pathology, Yale School of Medicine for the use of Tecan plate reader (Infinite 200 Pro). We also appreciate Caro Wiesehoefer for helping initial optimization of using the fluorescence plate reader, Gillian Clouser for helping mouse maintenance and antibody characterization, Mira Raju and Nathan Lang for helping initial characterization of waveform analysis of WT;*TG+* sperm, and Sofia Boeg Winge at department of growth and reproduction, Rigshospitalet, Copenhagen University Hospital for supporting human histology sample preparation. This study was supported by MCI Sokal award, NIH R01HD096745, Blavatnik Innovation Fund to J.J.C. J.Y.H. was a recipient of MCI postdoctoral fellowship.

## Author contributions

**Jae Yeon Hwang:** Conceptualization; methodology; formal analysis; data curation; writing - original draft; writing – review and editing

**Huafeng Wang:** Conceptualization; methodology; formal analysis; data curation

**Jong-Nam Oh;** Methodology; formal analysis; data curation

**Sarah F. Finnegan:** Methodology; formal analysis

**Niels E. Skakkebaek:** Methodology; formal analysis

**Jean-Ju Chung:** Conceptualization; formal analysis; data curation; writing - original draft; writing – review and editing; resource; supervision; funding acquisition

## References

Balbach, M., Rossetti, T., Ferreira, J., Ghanem, L., Ritagliati, C., Myers, R.W., et al. (2023). On-demand male contraception via acute inhibition of soluble adenylyl cyclase. Nature communications 14(1), 637.

Bearak, J., Popinchalk, A., Ganatra, B., Moller, A.-B., Tunçalp, Ö., Beavin, C., et al. (2020). Unintended pregnancy and abortion by income, region, and the legal status of abortion: estimates from a comprehensive model for 1990–2019. The Lancet Global Health 8(9), e1152–e1161.

Bharambe, N., Li, Z., Seiferth, D., Balakrishna, A.M., Biggin, P.C., and Basak, S. (2024). Cryo-EM structures of prokaryotic ligand-gated ion channel GLIC provide insights into gating in a lipid environment. Nature Communications 15(1), 2967.

Brown, S.G., Miller, M.R., Lishko, P.V., Lester, D.H., Publicover, S.J., Barratt, C.L., et al. (2018). Homozygous in-frame deletion in CATSPERE in a man producing spermatozoa with loss of CatSper function and compromised fertilizing capacity. Human Reproduction 33(10), 1812–1816.

Chang, Z., Qin, W., Zheng, H., Schegg, K., Han, L., Liu, X., et al. (2021). Triptonide is a reversible non-hormonal male contraceptive agent in mice and non-human primates. Nature communications 12(1), 1253.

Chen, S., Zhao, Y., Wang, Y., Shekhar, M., Tajkhorshid, E., and Gouaux, E. (2017). Activation and desensitization mechanism of AMPA receptor-TARP complex by cryo-EM. Cell 170(6), 1234–1246. e1214.

Chou, T.-H., Epstein, M., Fritzemeier, R.G., Akins, N.S., Paladugu, S., Ullman, E.Z., et al. (2024). Molecular mechanism of ligand gating and opening of NMDA receptor. Nature 632(8023), 209–217.

Chung, J.-J., Miki, K., Kim, D., Shim, S.-H., Shi, H.F., Hwang, J.Y., et al. (2017). CatSperζ regulates the structural continuity of sperm Ca2+ signaling domains and is required for normal fertility. Elife 6, e23082.

Chung, J.-J., Shim, S.-H., Everley, R.A., Gygi, S.P., Zhuang, X., and Clapham, D.E. (2014). Structurally distinct Ca2+ signaling domains of sperm flagella orchestrate tyrosine phosphorylation and motility. Cell 157(4), 808–822.

Chung, J.-J.L., Navarro, B., Krapivinsky, G., Krapivinsky, L., and Clapham, D.E. (2011). A novel gene required for male fertility and functional CATSPER channel formation in spermatozoa. Biophysical Journal 100(3), 90a.

Evans, R., O’Neill, M., Pritzel, A., Antropova, N., Senior, A., Green, T., et al. (2021). Protein complex prediction with AlphaFold-Multimer. biorxiv, 2021.2010.2004.463034.

Fagerberg, L., Hallström, B.M., Oksvold, P., Kampf, C., Djureinovic, D., Odeberg, J., et al. (2014). Analysis of the human tissue-specific expression by genome-wide integration of transcriptomics and antibody-based proteomics. Molecular & cellular proteomics 13(2), 397–406.

Gao, S., Yao, X., Chen, J., Huang, G., Fan, X., Xue, L., et al. (2023). Structural basis for human Cav1. 2 inhibition by multiple drugs and the neurotoxin calciseptine. Cell 186(24), 5363–5374. e5316.

Horvath, S., and Schreiber, C.A. (2017). Unintended pregnancy, induced abortion, and mental health. Current psychiatry reports 19, 1–6.

Huang, J., Pan, X., and Yan, N. (2024). Structural biology and molecular pharmacology of voltage-gated ion channels. Nature Reviews Molecular Cell Biology, 1–22.

Huang, X., Miyata, H., Wang, H., Mori, G., Iida-Norita, R., Ikawa, M., et al. (2023). A CUG-initiated CATSPERθ functions in the CatSper channel assembly and serves as a checkpoint for flagellar trafficking. Proceedings of the National Academy of Sciences 120(39), e2304409120.

Hwang, J.Y., and Chung, J.-J. (2023). CatSper calcium channels: 20 years on. Physiology 38(3), 125–140.

Hwang, J.Y., Mannowetz, N., Zhang, Y., Everley, R.A., Gygi, S.P., Bewersdorf, J., et al. (2019). Dual sensing of physiologic pH and calcium by EFCAB9 regulates sperm motility. Cell 177(6), 1480–1494. e1419.

Hwang, J.Y., Maziarz, J., Wagner, G.P., and Chung, J.-J. (2021a). Molecular evolution of CatSper in mammals and function of sperm hyperactivation in gray short-tailed opossum. Cells 10(5), 1047.

Hwang, J.Y., Nawaz, S., Choi, J., Wang, H., Hussain, S., Nawaz, M., et al. (2021b). Genetic defects in DNAH2 underlie male infertility with multiple morphological abnormalities of the sperm flagella in humans and mice. Frontiers in Cell and Developmental Biology 9, 662903.

Hwang, J.Y., Wang, H., Lu, Y., Ikawa, M., and Chung, J.J. (2022). C2cd6-encoded CatSpertau targets sperm calcium channel to Ca(2+) signaling domains in the flagellar membrane. Cell Rep 38(3), 110226. doi: 10.1016/j.celrep.2021.110226.

Ikawa, M., Nakanishi, T., Yamada, S., Wada, I., Kominami, K., Tanaka, H., et al. (2001). Calmegin is required for fertilin α/β heterodimerization and sperm fertility. Developmental biology 240(1), 254–261.

Lee, K.-H., and Hwang, J.Y. (2024). Ca2+ homeostasis and male fertility: a target for a new male contraceptive system. Animal Cells and Systems 28(1), 171–183.

Lin, S., Ke, M., Zhang, Y., Yan, Z., and Wu, J. (2021). Structure of a mammalian sperm cation channel complex. Nature 595(7869), 746–750.

Liu, J., Xia, J., Cho, K.-H., Clapham, D.E., and Ren, D. (2007). CatSperβ, a novel transmembrane protein in the CatSper channel complex. Journal of Biological Chemistry 282(26), 18945–18952.

Liu, X., and Wang, W. (2023). Asymmetric gating of a human hetero-pentameric glycine receptor. Nature communications 14(1), 6377.

Luque, G.M., Schiavi-Ehrenhaus, L.J., Jabloñski, M., Balestrini, P.A., Novero, A.G., Torres, N.I., et al. (2023). High-throughput screening method for discovering CatSper inhibitors using membrane depolarization caused by external calcium chelation and fluorescent cell barcoding. Frontiers in Cell and Developmental Biology 11, 1010306.

Mariani, N.A., Silva, J.V., Fardilha, M., and Silva, E.J. (2023). Advances in non-hormonal male contraception targeting sperm motility. Human Reproduction Update 29(5), 545–569.

Meng, E.C., Goddard, T.D., Pettersen, E.F., Couch, G.S., Pearson, Z.J., Morris, J.H., et al. (2023). UCSF ChimeraX: Tools for structure building and analysis. Protein Science 32(11), e4792.

Nadezhdin, K.D., Neuberger, A., Nikolaev, Y.A., Murphy, L.A., Gracheva, E.O., Bagriantsev, S.N., et al. (2021). Extracellular cap domain is an essential component of the TRPV1 gating mechanism. Nature communications 12(1), 2154.

Nand, K., Jordan, T.B., Yuan, X., Basore, D.A., Zagorevski, D., Clarke, C., et al. (2023). Bacterial production of recombinant contraceptive vaccine antigen from CatSper displayed on a human papilloma virus-like particle. Vaccine 41(46), 6791–6801.

Nickels, L., and Yan, W. (2024). Nonhormonal male contraceptive development—strategies for progress. Pharmacological Reviews 76(1), 37–48.

Nielsen, J.E., Rolland, A.D., Rajpert-De Meyts, E., Janfelt, C., Jørgensen, A., Winge, S.B., et al. (2019). Characterisation and localisation of the endocannabinoid system components in the adult human testis. Scientific reports 9(1), 12866.

Qi, H., Moran, M.M., Navarro, B., Chong, J.A., Krapivinsky, G., Krapivinsky, L., et al. (2007). All four CatSper ion channel proteins are required for male fertility and sperm cell hyperactivated motility. Proceedings of the National Academy of Sciences 104(4), 1219–1223.

Quill, T.A., Sugden, S.A., Rossi, K.L., Doolittle, L.K., Hammer, R.E., and Garbers, D.L. (2003). Hyperactivated sperm motility driven by CatSper2 is required for fertilization. Proceedings of the National Academy of Sciences 100(25), 14869–14874.

Ren, D., Navarro, B., Perez, G., Jackson, A.C., Hsu, S., Shi, Q., et al. (2001). A sperm ion channel required for sperm motility and male fertility. Nature 413(6856), 603–609.

Schindelin, J., Arganda-Carreras, I., Frise, E., Kaynig, V., Longair, M., Pietzsch, T., et al. (2012). Fiji: an open-source platform for biological-image analysis. Nature methods 9(7), 676–682.

Toyoda, Y., and Chang, M. (1974). Fertilization of rat eggs in vitro by epididymal spermatozoa and the development of eggs following transfer. Reproduction 36(1), 9–22.

Turney, T.S., Li, V., and Brohawn, S.G. (2022). Structural Basis for pH-gating of the K(+) channel TWIK1 at the selectivity filter. Nat Commun 13(1), 3232. doi: 10.1038/s41467-022-30853-z.

Wachten, D., Jikeli, J.F., and Kaupp, U.B. (2017). Sperm sensory signaling. Cold Spring Harbor perspectives in biology 9(7), a028225.

Wang, H., Dou, Q., Jeong, K.J., Choi, J., Gladyshev, V.N., and Chung, J.-J. (2022). Redox regulation by TXNRD3 during epididymal maturation underlies capacitation-associated mitochondrial activity and sperm motility in mice. Journal of Biological Chemistry 298(7).

Wang, H., McGoldrick, L.L., and Chung, J.-J. (2021). Sperm ion channels and transporters in male fertility and infertility. Nature Reviews Urology 18(1), 46–66.

Wang, J., Youkharibache, P., Zhang, D., Lanczycki, C.J., Geer, R.C., Madej, T., et al. (2020). iCn3D, a web-based 3D viewer for sharing 1D/2D/3D representations of biomolecular structures. Bioinformatics 36(1), 131–135.

Wang, L., Zhou, H., Zhang, M., Liu, W., Deng, T., Zhao, Q., et al. (2019). Structure and mechanogating of the mammalian tactile channel PIEZO2. Nature 573(7773), 225–229.

Watanabe, D., Okabe, M., Hamajima, N., Morita, T., Nishina, Y., and Nishimune, Y. (1995). Characterization of the testis-specific gene ‘calmegin’promoter sequence and its activity defined by transgenic mouse experiments. FEBS letters 368(3), 509–512.

Xiao, B. (2024). Mechanisms of mechanotransduction and physiological roles of PIEZO channels. Nature Reviews Molecular Cell Biology, 1–18.

Young, S., Schiffer, C., Wagner, A., Patz, J., Potapenko, A., Herrmann, L., et al. (2024). Human fertilization in vivo and in vitro requires the CatSper channel to initiate sperm hyperactivation. The Journal of Clinical Investigation 134(1).

Yue, F., Cheng, Y., Breschi, A., Vierstra, J., Wu, W., Ryba, T., et al. (2014). A comparative encyclopedia of DNA elements in the mouse genome. Nature 515(7527), 355–364.

Zhao, Y., Wang, H., Wiesehoefer, C., Shah, N.B., Reetz, E., Hwang, J.Y., et al. (2022). 3D structure and in situ arrangements of CatSper channel in the sperm flagellum. Nature communications 13(1), 3439.

Zhu, S., Noviello, C.M., Teng, J., Walsh Jr, R.M., Kim, J.J., and Hibbs, R.E. (2018). Structure of a human synaptic GABAA receptor. Nature 559(7712), 67–72.

