## Supplementary Information for "CATSPERε extracellular domains are essential for sperm calcium channel assembly and activity modulation"

**List of materials**

Figure S1. *CATSPER* gene expression in mouse and human, Related to Figure 1.

Figure S2. Generation of the *Catspere*-null mice by CRISPR/Cas9 genome editing, Related to Figure 1 and 2.

Figure S3. Unaltered hyperactivation and protein levels of CATSPER subunits in spermatozoa from WT mice expressing the transgene encoding ECDs-truncated CATSPERε, Related to Figure 3.

Figure S4. No change in the impaired sperm hyperactivation and male infertility of *Catspere*-null mice carrying the transgene encoding truncated CATSPERε, Related to Figure 5.

Figure S5. Predicted impact of individual CATSPERε ECD domain on the canopy structure formation and solubility of serially deleted recombinant CATSPERε ECD proteins, Related to Figure 6.

### Supplementary Figures and Legends

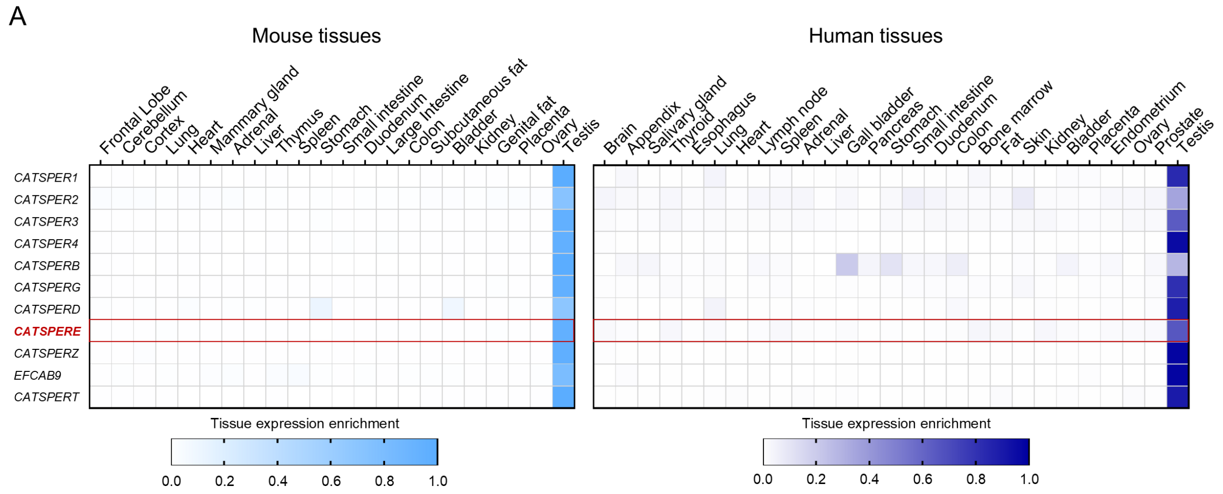


B

```
Mouse_P0DP43    MPSAGQRKPGSLLALQALQKWLLRGGVGAMLARQVVAALLLWLSCCVSALWRYYINSQDYSIFSTRSSIKLEYEGNSFVSWKIPESCKVENTTSPKTTLHCKRAGIHTIKPIA-GNQEVE   119
Human_Q5SY80    ------------------------------MSAREVAVLLLWLSCYGSALWRYSTNSPNYRIFSTRSTIKLEYEGTLFTEWSVPETCFVLNKSSPTTELRCSSPGVHAIKPIVTGPDEEE   90
                                              :: : **.*******  ******  ** :* ******:*******. *..*.:**:* * *.:**.* *:*.  *:*:****. * :* *

Mouse_P0DP43    RHLTVDNSYICYLWYFTVVDVYYNLSQIVTIWVYDPESASTEELIWTAKKPSLSSRVLTKQMNTLGQRPFIFTVEKRLTYHPGPLTSEGTWVIHLPMSSDDIAKVIRGNKVAFQDCFIAN   239
Human_Q5SY80    RYLFVESSHTCFLWYYRVRHFFNNFTQLITVWAYDPESADPDELLGNAEEPSINSIVLSTQMATLGQKPVIHTVLKRKVYSSNEKMRRGTWRIVVPMTKDDALKEIRGNQVTFQDCFIAD   210
                *:* *:.*: *:***: * ..: *::*::*:*.******. :**: .*::**:.* **:.** ****:*.*.** ** .*   .    .*** * :**:.**  * ****:*:*******:

Mouse_P0DP43    LYFMLTYPMTIISEPPGYEPLTVPPGSPLMLSWDTCISTFALLATDQETFQTNDSFQTWTRVRAPPGILSDAQRHSLRDVIIFDQGTLFLVDGTVYLRTEDEFTKLDESRGISETGILGF   359
Human_Q5SY80    FLILLTFPLLTIPEIPGYLPISSPRGSQLMASWDACVVASAVLVTDMETFHTTDSFKSWTRIRVPPDILSDDERRSVAHVILSRDGIVFLINGVLYIKSFRGFIRLGGIVNLPDGGITGI   330
                : ::**:*:  * * *** *:: * ** ** ***:*: : *:*.** ***:*.***::***:*.**.**** :*:*: .**:  :* :**::*.:*:::   * :*.   .: : ** *:

Mouse_P0DP43    SKRRWCQIRYLYKLASKKSILIAWSKTTVYAGYATFRFVTLTDTAKLKDFLKLPQTDTLEVMSVEYLWHPLEAAVLLSHCSVCTTNTRNIRIVIYSAIFQTWTLQDFELQLPKEAILEFR   479
Human_Q5SY80    SSRKWCWVNYLLKAKGRRSTFAVWTENEIYLGSILLKFARLVTTTELKNILSLSVTATLTIDRVEYTGHPLEIAVFLNYCTVCNVT-KKIFLVIYNEDTKQWVSQDFTLDAPIDSVTMPH   449
                *.*:** :.** *  .::* : .*::. :* *   ::*. *. *::**::*.*  * ** :  ***  **** **:*.:*:**... ::* :***.   : *. *** *: * :::   :

Mouse_P0DP43    FLYSAMPDIIMWDQHHVYYSYKNFTVVGTISTPSGETNLSSLSQGSKIHQVLTDRIGNVVVKMENNVMFYIKADITEAVILHTWVNTTAKTVVLFDKSFEVCILYYNENLDEKYQLQTQP   599
Human_Q5SY80    FTFSALPGLLLWNKHSIYYCYHNFTFTGILQTPAGHGNLSMLSNDSIIHEVFIDYYGDILVKMENNVMFYIKADITEAVILHTWVNTTAKTVVLFDKSFEVCILYYNENLDEKYQLQTQP   565
                * :**:*.:::*::* :**.*:***..* :.**:*. *** **:.* **:*: *  *:::*******:** * :  :** ** *.* *:::.:::..* :. :**      :.  :* *

Mouse_P0DP43    YPLILELQSINKDLGDWCPYLAFQHNIHSQFYHMDKGESLTIWSQIVYPENRGLYIVVEHYGSSVMTWTQNLEYEIASGFCTKTMITRFFQTTNYELVDNYYQLQKENTGLMLLQFRPSE   719
Human_Q5SY80    YPLHLEAQSIAFTTKDKCPYMAFHNNVAHVFYFLDKGEALTVWTQIVYPENTGLYVIVESYGPKILQESHEISFEAAFGYCTKTLTLTFYQNVDYERISDYFETQDKHTGLVLVQFRPSE   685
                *** ** ***     * ***:**::*:   **.:****:**:*:******* ***::** ** .::  ::::.:* * *:****:   *:*..:** :.:*:: *.::***:*:******

Mouse_P0DP43    FSRTCLTAKPVFEIDVGCDSSKYIMVRGFNKSRCQRRDFSYVIDKELLRESLSDNLKVRYDVAKYGCPLTLELGQMFQPIVELYDENGFIKIVDANFILWEIHGRNDYTFNSTMEQNGCI   839
Human_Q5SY80    YSKACPIAQKVFQIAVGCDDKKFIAIKGFSKKGCHHHDFSYVIEKSYLRHQPSKNLRVRYIWGEYGCPLRLDFTEKFQPVVQLFDDNGYVKDVEANFIVWEIHGRDDYSFNNTMAQSGCL   805
                :*::*  *: **:* ****..*:* ::**.*. *:::******:*. **.. *.**:***  .:***** *:: : ***:*:*:*:**::* *:****:******:**:**.** *.**:

Mouse_P0DP43    NEAQTWDSMIEENPDIPLDDVWGPQNYRPCFSYAIGKPGDLGQPYEILNYSNKNHIKWPMTYAGMYVYRLKILDPNYSFCNLTTIFAIESLGMIPRSSVYLVAALIFVLMLTFISILVLS   959
Human_Q5SY80    HEAQTWKSMIELNKHLPLEEVWGPENYKHCFSYAIGKPGDLNQPYEIINSSNGNHIFWPMGHSGMYVFRVKILDPNYSFCNLTAMFAIETFGLIPSPSVYLVASFLFVLMLLFFTILVLS   925
                :*****.**** * .:**::****:**: ***********.*****:* ** *** *** ::****:*:************::****::*:**  ******::***** *::*****

Mouse_P0DP43    YFWYLKIYRQFIIEPLHKRPAKQKKN   985
Human_Q5SY80    YFRYMRIYRRYIYEPLHKPQRKRKKN   951
                ** *::***::* *****   *:***
```

**Figure S1. *CATSPER* gene expression in mouse and human, Related to Figure 1**. (A) Tissue enrichment of the transcripts encoding CATSPERε and other CATSPER subunits. Shown are heatmaps for the enriched transcript levels of *CATSPER1*, *2*, *3*, *4*, *B*, *G*, *D*, *E*, *Z*, *T*, and *EFCAB9* in various tissues of mouse (*left*) and human (*right*). Enrichment levels for each subunit are calculated by normalizing transcript levels in each tissue by total amount from the entire tissues in comparison. (B) Amino acid sequence alignment of mouse and human CATSPERε. Highlighted in yellow are the epitope sequences for human and mouse CATSPERε antibodies (α-hε-31 and α-mε-331, respectively) used in this study.

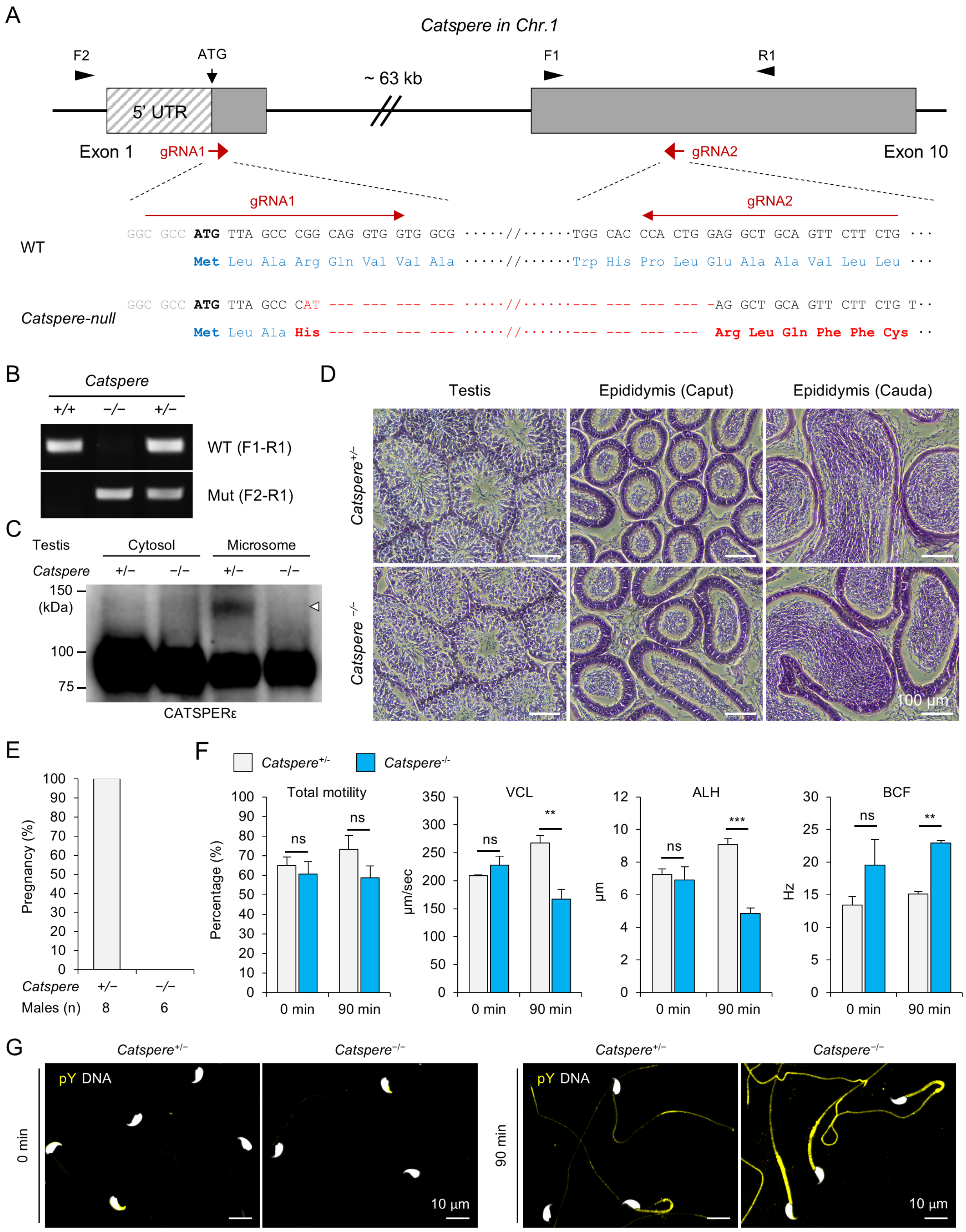
A
Catspere in Chr.1
F2
ATG
~ 63 kb
F1
R1
5' UTR
Exon 1
gRNA1
gRNA2
Exon 10
WT
GGC GCC ATG TTA GCC CGG CAG GTG GTG GCG ·····//······TGG CAC CCA CTG GAG GCT GCA GTT CTT CTG ···
Met Leu Ala Arg Gln Val Val Ala ·····//······Trp His Pro Leu Glu Ala Ala Val Leu Leu ···
Catspere-null
GGC GCC ATG TTA GCC CAT --- --- --- --- ·····//·········--- --- --- --- -AG GCT GCA GTT CTT CTG T··
Met Leu Ala His --- --- --- --- ·····//·········--- --- --- --- Arg Leu Gln Phe Phe Cys ··
B
Catspere
+/+ −/− +/−
WT (F1-R1)
Mut (F2-R1)
C
Testis
Cytosol
Microsome
Catspere +/− −/− +/− −/−
150
(kDa)
100
75
CATSPERε
D
Testis
Epididymis (Caput)
Epididymis (Cauda)
Catspere+/−
Catspere −/−
100 µm
E
Pregnancy (%)
100 90 80 70 60 50 40 30 20 10 0
Catspere +/− −/−
Males (n) 8 6
F
Catspere+/-
Catspere-/-
Total motility
Percentage (%)
VCL
µm/sec
ALH
µm
BCF
Hz
ns
**
***
0 min
90 min
G
Catspere+/−
Catspere−/−
pY DNA
0 min
90 min
10 µm

**Figure S2. Generation of the *Catspere*-null mice by CRISPR/Cas9 genome editing, Related to Figure 1 and 2.** (A) *Catspere*-null allele with ~63 Kb deletion of genomic region by CRISPR/Cas9. Marked in red are non-native amino acids resulted from frameshift in *Catspere*-null allele. Arrows indicate the locations of genotyping primers. (B) Genotyping of *Catspere*-null mice by genomic DNA PCR with the primers marked in panel A. (C) Immunoblotting of CATSPERε in the cytosol and microsome fractions of testis from heterozygous (+/-) and homozygous (-/-) *Catspere*-mutant mice. A white arrowhead indicates CATSPERε. (D) Histology of *Catspere*-null testis and epididymis. Shown are hematoxylin and eosin (H&E)-stained sections of testis (*left*), and caput (*middle*) and cauda (*right*) epididymis from *Catspere*$^{+/-}$ (*top*) and *Catspere*$^{-/-}$ (*bottom*) males. (E) Pregnancy rate of WT females mated with *Catspere*$^{+/-}$ and *Catspere*$^{-/-}$ males. (F) Computer Assisted Semen Analysis (CASA) for *Catspere*$^{+/-}$ (gray) and *Catspere*$^{-/-}$ (blue) sperm motility. Total sperm motility and hyperactivation parameters such as curvilinear velocity (VCL) and amplitude of lateral head (ALH), as well as beat cross frequency (BCF) are measured. ns, non-significant, **p<0.01, and ***p<0.01. N=3. Data are represented as mean ± SEM. (G) Confocal images for immunostained protein tyrosine phosphorylation in *Catspere*$^{+/-}$ and *Catspere*$^{-/-}$ sperm before (0 min) and after (90 min) inducing capacitation. Hoechst is used for DNA staining.

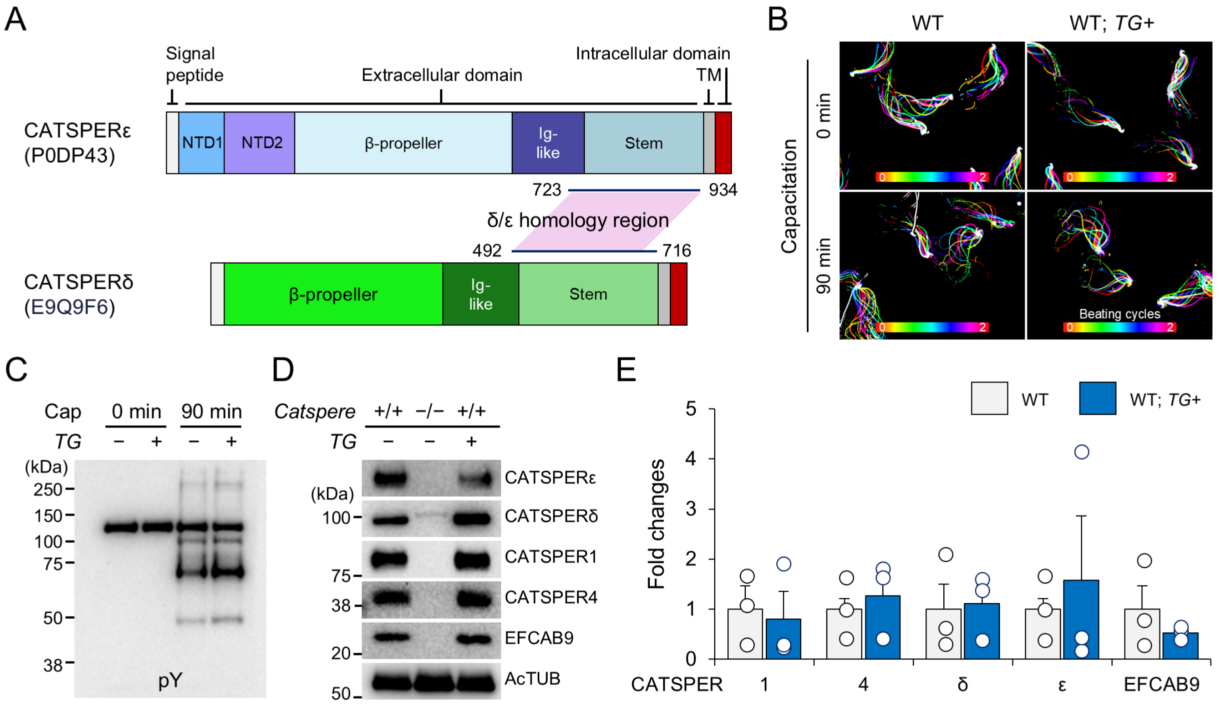


**Figure S3. Unaltered hyperactivation and protein levels of CATSPER subunits in spermatozoa from WT mice expressing the transgene encoding ECDs-truncated CATSPERε, Related to Figure 3.** (A) Comparison of mouse CATSPERε and CATSPERδ domain organization. Schematic diagram is mouse CATSPERε (Uniprot: P0DP43) and CATSPERδ (Uniprot: E9Q9F6) marked with C-terminal region of sequence homology. (B) Flagellar waveforms of WT (*left*) and WT with the transgene (WT; *TG+, right*) before (0 min, *top*) and after (90 min, *bottom*) inducing capacitation. Shown are the overlaid images of sperm flagella for 2 beating cycles. Time information is color-coded. (C) Immunoblotting of protein tyrosine phosphorylation (pY) in WT and WT; *TG+* sperm. pY development was examined in sperm before (0 min) and after (90 min) inducing capacitation (Cap). (D) Protein levels of CATSPER subunits in WT and WT; *TG+* sperm by immunoblotting. *Catspere*-null sperm were used as negative control. Acetylated tubulin (AcTUB) is loading control. (E) Fold changes of the CATSPER subunits in WT; *TG+* sperm (blue) compared to WT sperm (gray). Circles indicate relative protein levels in sperm from each male. Average levels of CATSPER subunits in WT sperm are set to 1. Levels of CATSPER subunits in WT; *TG+* sperm are not significantly different from those in WT sperm. Band intensities from independent immunoblotting were measured to quantify protein levels. Data are represented as mean ± SEM. N=3.

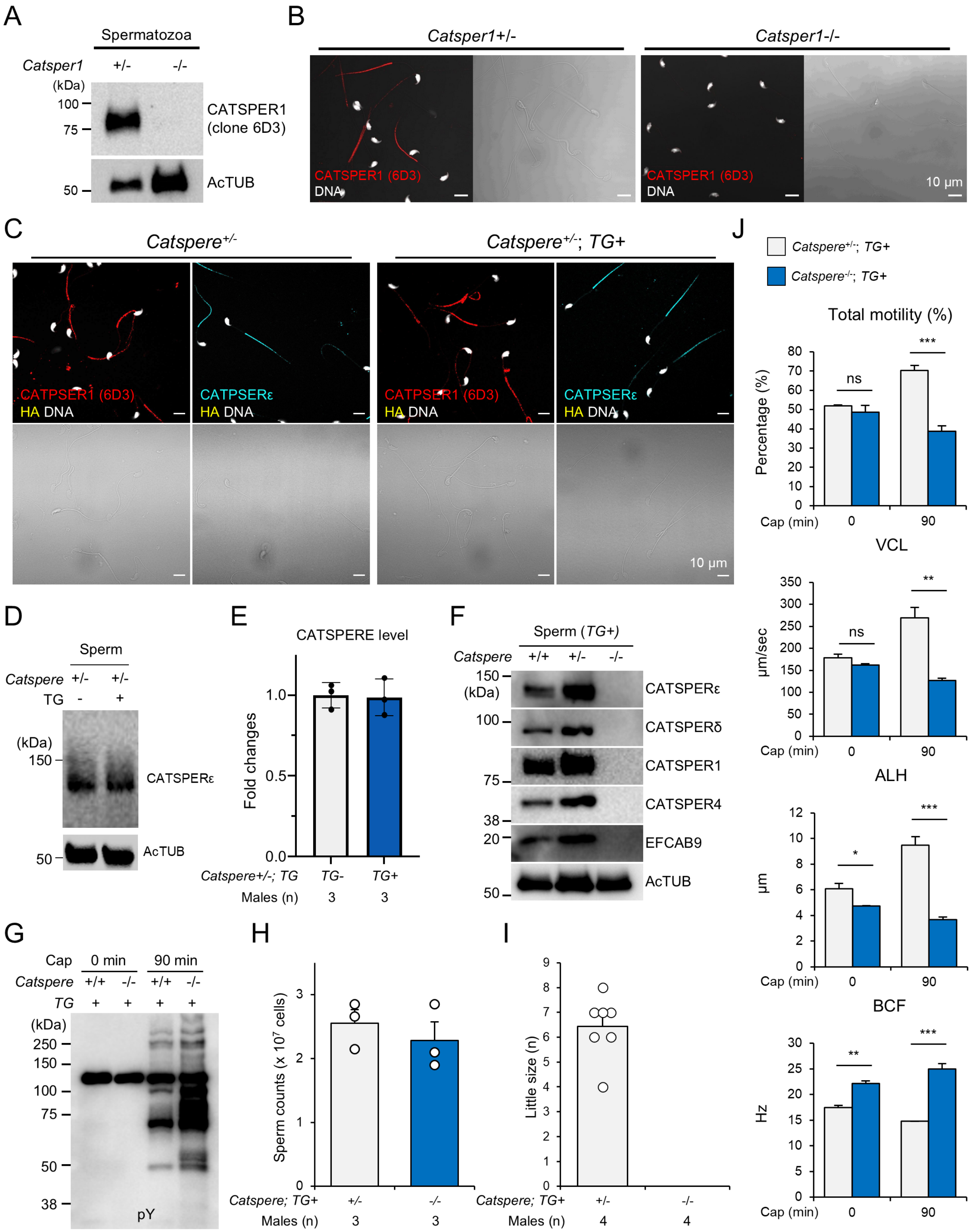

A
Spermatozoa
Catsper1 +/- -/-
(kDa)
100
75
50
CATSPER1 (clone 6D3)
AcTUB
B
Catsper1+/-
Catsper1-/-
CATSPER1 (6D3)
DNA
10 μm
C
Catspere+/-
Catspere+/-; TG+
CATPSER1 (6D3)
HA DNA
CATPSERε
10 μm
D
Sperm
Catspere +/- +/-
TG - +
(kDa)
150
CATSPERε
50
AcTUB
E
CATSPERE level
Fold changes
1.0
0.5
0.0
Catspere+/-; TG TG- TG+
Males (n) 3 3
F
Sperm (TG+)
Catspere +/+ +/- -/-
150
(kDa)
100
75
38
20
50
CATSPERε
CATSPERδ
CATSPER1
CATSPER4
EFCAB9
AcTUB
G
Cap 0 min 90 min
Catspere +/+ -/- +/+ -/-
TG + + + +
(kDa)
250
150
100
75
50
38
pY
H
Sperm counts (x 10^7 cells)
Catspere; TG+ +/- -/-
Males (n) 3 3
I
Little size (n)
Catspere; TG+ +/- -/-
Males (n) 4 4
J
Catspere+/-; TG+
Catspere-/-; TG+
Total motility (%)
Percentage (%)
ns
***
Cap (min) 0 90
VCL
μm/sec
ns
**
ALH
μm
*
***
BCF
Hz
**
***

**Figure S4. No change in the impaired sperm hyperactivation and male infertility of the *Catspere*-null mice carrying the transgene encoding truncated CATSPERε, Related to Figure 5**. (A-B) The specificity of monoclonal anti-CATSPER1 (clone 6D3) was established in this study. Immunoblotting of whole sperm lysate (A) and immunofluorescence (*left*) and corresponding DIC (*right*) images from confocal microscopy (B). Hoechst counterstains DNA. *Catsper1+/-* and *Catsper1-/-* sperm were used for positive and negative control, respectively. (C) Confocal images of sperm from *Catspere*$^{+/-}$ males with (*right*) or without (*left*) the transgene encoding truncated CATSPERε (*TG*). Shown are fluorescence (*top*) and corresponding DIC (*bottom*) images. Truncated CATSPERε (yellow) and CATSPER1 (red) or native full-length CATSPERε (cyan) were co-immunostained. Truncated CATSPERε was probed against HA. Hoechst counterstains DNA. (D) Immunoblotting of the native CATSPERε in *Catspere*$^{+/-}$ epididymal sperm with or without TG. Acetylated tubulin (AcTUB) is a loading control. (E) Quantitative comparison of CATSPERε levels in *Catspere*$^{+/-}$ (gray bars) and *Catspere*$^{+/-}$; *TG*+ (blue bars) epididymal sperm. Circles indicate the protein levels in sperm from individual males. CATSPERε levels in *Catspere*$^{+/-}$ sperm are set to 1-fold. Data are represented as mean ± SEM. N=3. (F) Protein levels of CATSPER subunits in sperm from WT (+/+) and *Catspere*-heterozygous (+/-) or knockout (-/-) mice expressing the transgene (*TG*) encoding truncated CATSPERε. Acetylated tubulin (AcTUB) is a loading control. (G) Protein tyrosine phosphorylation of sperm from WT (+/+) and *Catspere*-null (-/-) mice expressing the transgene (*TG*) before (0 min) and after (90 min) inducing capacitation (Cap). (H) Counts for epididymis sperm from *Catspere*-heterozygous (*Catspere*+/-; *TG*+, gray, N=3, 2.55 ± 0.21 x $10^7$ cells) and knockout (*Catspere*-/-; *TG*+, blue bar, N=3, 2.28 ± 0.30 x $10^7$ cells) males carrying the transgene. Circles indicate sperm counts from individual males. (I) Litter size from pregnant females mated with *Catspere+/-*; *TG*+ or *Catspere-/-*; *TG*+ males. Circles represent the number of pups from each litter. (J) Computer-assisted semen analysis (CASA) to analyze *Catspere*$^{+/-}$; *TG*+ (gray) and *Catspere*$^{-/-}$; *TG*+ (blue) sperm motility. Total motility, curvilinear velocity (VCL), amplitude of lateral head (ALH), and beat cross frequency (BCF) were measured. ns, non-significant, *p<0.01, **p<0.01, and ***p<0.01. N=3. Data are represented as mean ± SEM (E, H, I, and J).

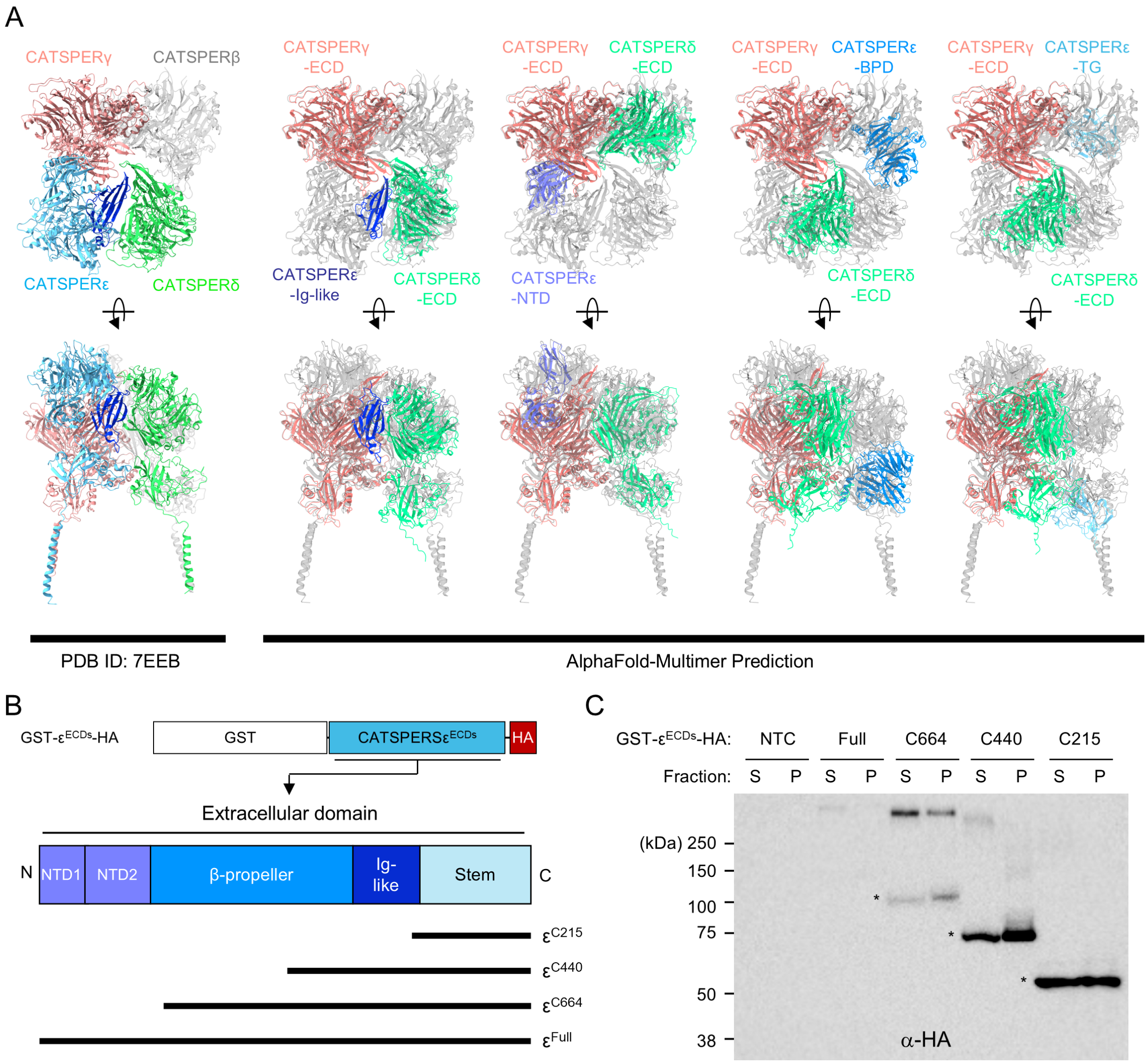


**Figure S5. Predicted impact of individual CATSPERε ECD domain on the canopy structure and solubility of serially deleted recombinant CATSPERε ECD proteins, Related to Figure 6.** (A) Predicted interaction of individual extracellular domains (ECDs) of CATSPERε with those of CATSPERγ and CATSPERδ. Shown are topdown (*top*) and side (*bottom*) views of the reported CATSPER canopy structure (*left*; PDB: 7EEB) and AlphaFold multimer-predicted heterotrimers (*right*) of each ECD domains for CATSPERε - Ig-like domain, N-terminal domain (NTD), or β-propeller domain (BPD) - or ECD regions of the truncated CATSPERε expressed from the transgene (TG) with CATSPERγ and CATSPERδ ECDs. The modeled trimer structures are fitted to the reference canopy structure (gray transparent). CATSPERε Ig-like domain is predicted to form a trimer only with ECDs of CATSPERγ and CATSPERδ, which fits well into the reference structure (gray). (B-C) Serially truncated recombinant CATSPERε ECD proteins and their solubility. (B) Schematic diagram of fragments of recombinant CATSPERε ECDs tagged with GST and HA at the N- and C-terminus, respectively, designed to test for their partitioning in soluble vs.

pellet fractions. The fragments are designed by serial truncation of full-length ECD ($\varepsilon^{Full}$) from N-terminus, which are 215 ($\varepsilon^{C215}$), 440 ($\varepsilon^{C440}$), and 664 ($\varepsilon^{C664}$) amino acids length. (C) Recombinant CATSPERε ECDs are expressed in 293T cells, and the proteins were detected from soluble (S) and insoluble pellet (P) fractions. Recombinant proteins are probed by HA tag. Asterisks (*) indicate predicted sizes of the recombinant proteins.
